# Multimodal evidence for impaired glymphatic brain clearance as a mediator of “inflammaging” in schizophrenia – integrating insights from in vivo neuroimaging, omics, and post-mortem single nucleus-RNA-Seq data

**DOI:** 10.64898/2026.09.24.753662

**Authors:** Xiaolin Tan, Malte Simon, Daniel Rosebrock, Mariia Firuleva, Doris Gruber-Schoffnegger, Marisa Fuerst, Rana Aldisi, Stefanie Heilmann-Heimbach, Karl Bechter, Gereon Schnellbaecher, Thomas Frodl, Klaus Mathiak, Arnim Johannes Gaebler

**Affiliations:** Shengjing Hospital of China Medical University, Shenyang, China; Evotec SE, Hamburg, Germany; Institute for Genomic Statistics and Bioinformatics, University of Bonn, Bonn, Germany; Institute of Human Genetics, University of Bonn, School of Medicine & University Hospital Bonn, Bonn, Germany; Department of Psychiatry and Psychotherapy II, Ulm University, Germany; Department of Psychiatry, Psychotherapy and Psychosomatics, Faculty of Medicine, RWTH Aachen University, Aachen Germany

## Abstract

Persistent cognitive impairment and reduced life expectancy in schizophrenia may arise from inflammation-driven accelerated aging (“inflammaging”), yet the underlying mechanisms remain unclear. Impaired glymphatic clearance, dependent on polarized aquaporin-4 (AQP4) localization at astrocytic endfeet, offers a plausible mechanistic link between inflammation and brain aging.

Here, we integrate in vivo neuroimaging with post-mortem single-nucleus transcriptomics from distinct patient cohorts to demonstrate glymphatic dysfunction in schizophrenia and define its molecular basis. We identify reduced astrocytic expression of components of the dystrophin-associated protein complex (SNTA1, DAG1) required for AQP4 polarization. Both in vivo neuroimaging and post-mortem gene expression signatures of glymphatic dysfunction are restricted to an inflammatory subgroup characterized by peripheral immune activation and pro-inflammatory glial transcriptional profiles. Moreover, imaging signatures of glymphatic impairment are associated with schizophrenia polygenic risk scores enriched for immune- and oxidative stress-related aging pathways as well as cognitive dysfunction. At the cellular level, glymphatic dysfunction coincides with loss of homeostatic programs in microglia and astrocytes, increased inflammatory signaling, and endothelial signatures of blood– brain barrier disruption. Across modalities, glymphatic impairment is linked to accelerated aging, including increased brain and epigenetic age, and aging-related gene expression signatures.

**Graphical abstract:** 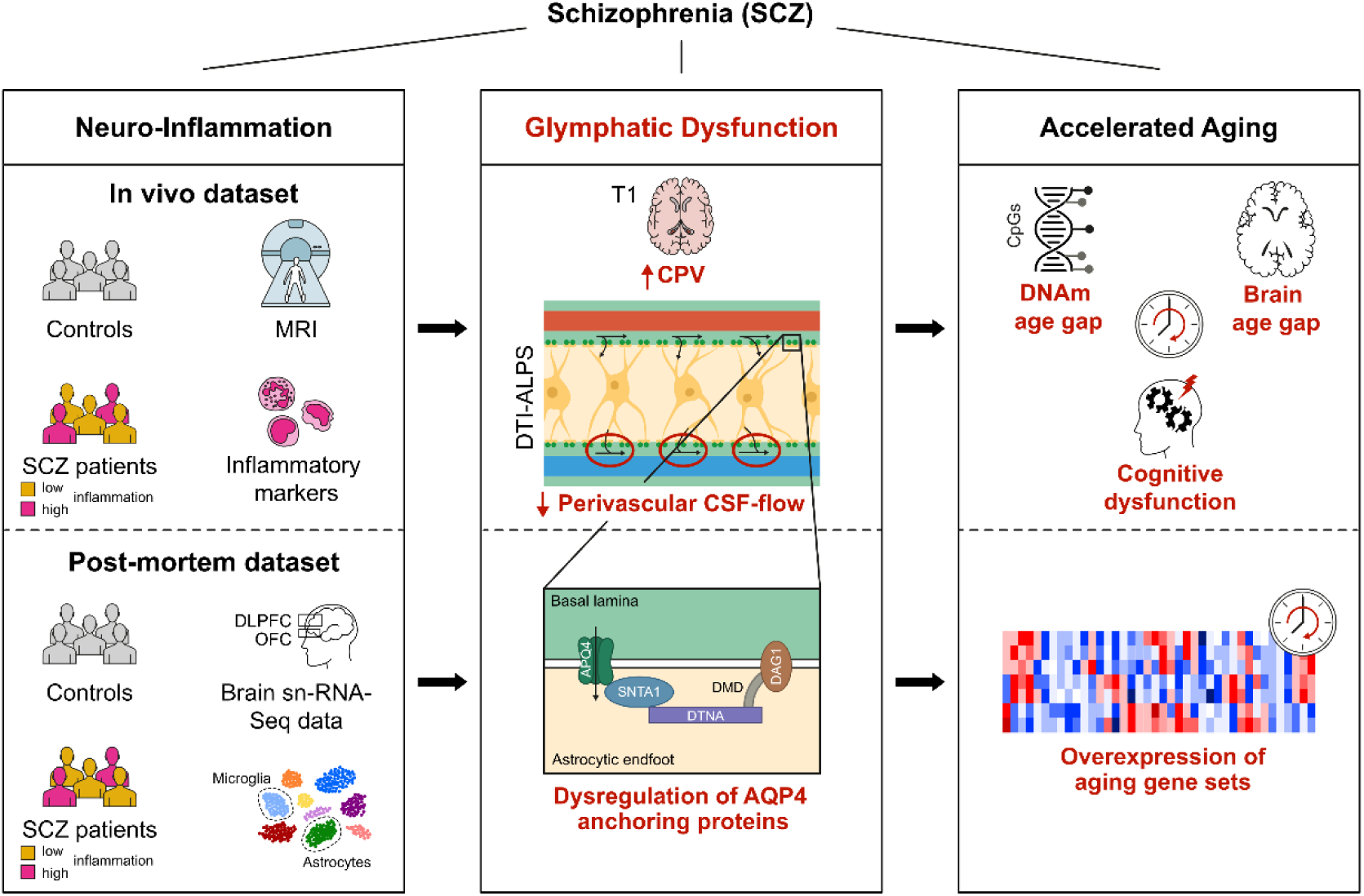

## 1. Introduction

Schizophrenia (SCZ) is a severe mental illness characterized by disturbances in perception, emotion, and cognition. Among the most pressing clinical challenges are persistent cognitive deficits and a markedly reduced life expectancy of approximately 15 years [1]. Cognitive impairment, a defining feature since Kraepelin’s original description of “dementia praecox,” remains a key determinant of long-term functional outcome in a substantial proportion of patients [2]. Although the underlying biology is incompletely understood, current models integrate early neurodevelopmental alterations with progressive neurobiological or neurodegenerative changes over the course of illness [3, 4].

Consistent with this framework, neuroimaging studies have demonstrated subtle but progressive brain volume loss associated with poorer clinical outcomes, while machine learning approaches have identified patterns indicative of accelerated brain aging [5]. Clinically, a subgroup of patients shows a deteriorating course marked by progressive cognitive decline, reminiscent of Kraepelin’s original concept and exhibiting biological overlap with frontotemporal dementia [6]. Beyond the central nervous system, accelerated aging signatures—including telomere shortening [7, 8], epigenetic age acceleration [9, 10], mitochondrial dysfunction [11], increased oxidative stress [12] and inflammatory signaling [13] - Together this findings suggest a systemic process that may contribute to the excess mortality observed in schizophrenia beyond suicide alone. Inflammation has emerged as a central mechanism linking to accelerated aging and cognitive dysfunction in SCZ. Elevated peripheral inflammatory markers are consistently observed in subsets of patients with more severe cognitive deficits or treatment-resistant symptoms [14]. Converging evidence from post-mortem studies and transcriptomic analysis indicates inflammatory brain signatures in a subgroup of patients, characterized by altered gene expression in microglia and astrocytes, dysregulation of cytokines and immune signaling pathways, and infiltration of peripheral immune cells [15–20]. These data reflect a shift from homeostatic to reactive glial states.

A potential pathway connecting inflammation, brain aging, and cognitive decline is dysfunction of the glymphatic system, a perivascular network responsible for the clearance of interstitial waste [21]. This system facilitates the influx of cerebrospinal fluid (CSF) along periarterial spaces, its exchange with interstitial fluid via aquaporin-4 (AQP4) channels localized to astrocytic endfeet, and subsequent drainage along perivenous pathways into meningeal lymphatic vessels. Impaired glymphatic function has been implicated in neurodegenerative disorders through reduced clearance of neurotoxic proteins such as amyloid-β and is sensitive to inflammatory processes in experimental models [22].

In humans, glymphatic function can be assessed non-invasively using indirect imaging markers, including the diffusion tensor imaging along perivascular space (DTI-ALPS) index [23] and choroid plexus volume (CPV) [24]. Emerging studies report reduced ALPS index and increased CPV in schizophrenia [25, 26]. However, the mechanistic underpinnings and clinical relevance of these abnormalities remain unclear.

Here, we address this gap by integrating multimodal in vivo imaging and multi-omics data with post-mortem single-nucleus transcriptomics across distinct patient cohorts. We test the hypothesis that glymphatic dysfunction—reflected by reduced ALPS index and increased CPV—is associated with peripheral inflammation, genetic risk enriched for immune and aging pathways, and signatures of accelerated biological aging, including brain and epigenetic age gaps, as well as cognitive impairment. In post-mortem tissue, we further examine molecular alterations in astrocytic endfoot organization, including AQP4 anchoring complex components and their relationship to inflammatory, vascular, and aging-associated transcriptional programs across cell types. By integrating these complementary approaches, we aim to provide a mechanistically informed characterization of glymphatic dysfunction in schizophrenia and its relevance for accelerated aging and cognitive decline.

## 2. Results

### 2.1 Reduced DTI-ALPS and enlarged CPV in schizophrenia are interconnected and independent from antipsychotic medication

Both the DTI-ALPS index and CPV are considered MRI-based markers related to glymphatic system function (Figure 1A). In this context, the DTI-ALPS index estimates water diffusion along perivascular spaces, which is thought to reflect CSF movement as part of the glymphatic clearance route. In different neurological conditions (e.g. Alzheimer’s disease, Parkinson’s disease), reduced ALPS index co-occurred with enlarged CPV suggesting it as a further indirect imaging marker of glymphatic dysfunction [27, 28]. Although, recent studies have demonstrated both reduced DTI-ALPS index and enlarged CPV in patients with SCZ, the relationship between the markers in SCZ, as well as their potential dependence on variables such as antipsychotic medication remain unclear.

**Figure 1.**
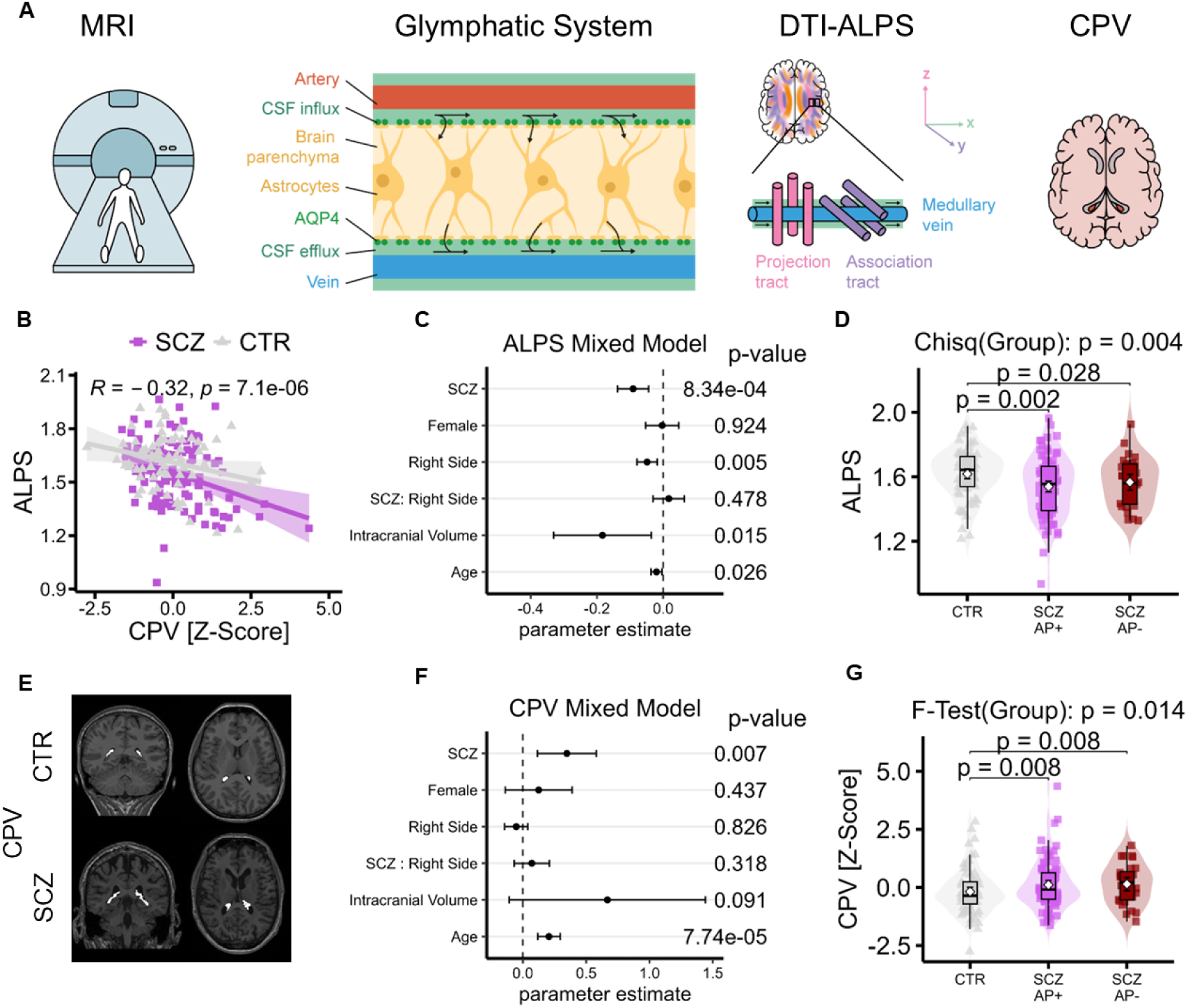
A. Illustration of the in vivo MRI study. Glymphatic clearance involves CSF flow along perivascular spaces (second panel). The DTI-ALPS index is used as an indirect measure of this process by quantifying water diffusion along the transversal axis (x-axis) in a paraventricular region of interest, where major white matter tracts that would otherwise dominate the diffusion signal run predominantly perpendicular to this axis. Choroid plexus volume (CPV) was used as a complementary indirect measure of glymphatic clearance. B. A negative correlation between DTI-ALPS index and CPV emerged for both patients with SCZ and CTR subjects, i.e. lower DTI-ALPS index was associated with higher CPV. C. The DTI-ALPS index was significantly reduced in SCZ. Across both groups, lower values were also observed for the right as compared to the left hemisphere, subjects with higher intracranial volume or higher age. No lateralization differences emerged between groups. 90% confidence intervals were used for variables tested with one-sided (right-tailed) hypotheses (group, gender, laterality, age), whereas 95% confidence intervals were used for variables tested with two-sided hypotheses. D. Both patients receiving antipsychotic treatment (SCZ AP+, N = 87) and antipsychotic-free patients (SCZ AP-, N = 28) exhibited significantly lower DTI-ALPS than healthy control subjects. Inferential statistics are based on a linear model employing age, total intracranial volume and gender as further independent variables. To account for unequal variances, the linear model was estimated with HC3 robust standard errors. Overall group differences were assessed using Wald’s chi-square test. Estimated marginal means contrasts were used to compare the two SCZ subgroups with the CTR group, and false discovery rate (FDR) correction was applied to control for multiple comparisons. Error bars represent confidence intervals. E. Example of choroid plexus segmentation in a patient with SCZ and a CTR subject F. Patients with SCZ showed significantly higher CPV than healthy controls. Moreover, a significant effect of age emerged, indicating higher age is associated with higher CPV. 90% confidence intervals were used for variables tested with one-sided (right-tailed) hypotheses (group, gender, laterality, age), whereas 95% confidence intervals were used for variables tested with two-sided hypotheses. G. Both medicated (SCZ AP+, N = 87) and antipsychotic-free patients (SCZ AP-, N = 28) exhibited significantly higher CPV than CTR subjects, excluding pure medication effects. Since the outcome variables contained outliers, robust linear models were fitted. Overall group differences were assessed using Wald-type F tests. Estimated marginal means contrasts were used to compare the two SCZ subgroups with the CTR group, and false discovery rate (FDR) correction was applied to control for multiple comparisons. Error bars represent confidence intervals.

We studied DTI-ALPS and CPV in a cohort of 115 patients with SCZ and 73 control subjects. As expected, CPV demonstrated significant negative correlation with DTI-ALPS index among the whole sample (t(183) = −3.787, p < 0.001) (Figure 1B). For a systematic comparison of DTI-ALPS index and CPV between patients with SCZ and healthy control subjects we employed linear mixed models considering further influential variables including gender, laterality, the total intracranial volume and age (Figure 1C). As compared to healthy control subjects, patients with SCZ exhibited significantly lower DTI-ALPS index and higher CPV. As expected, both DTI-ALPS index and CPV showed a significant association with age indicating age-related impairment of glymphatic clearance across the whole sample. Both groups showed higher DTI-ALPS index in the left compared with the right hemisphere, whereas no significant lateralization pattern emerged for CPV. No significant differences in lateralization were observed between groups. Moreover, the DTI-ALPS index was negatively correlated with the total intracranial volume. Although CPV was already normalized to the total intracranial volume, a positive correlation emerged – at a trend level – too. No significant gender differences emerged. To rule out medication effects, we performed a subgroup analysis comparing patients with and without antipsychotic treatment. Importantly, unmedicated patients showed similar findings to medicated patients, exhibiting a significantly lower DTI-ALPS index (Figure 1D) and higher CPV (Figure 1G) than healthy controls, suggesting disease-related rather than medication-related effects.

### 2.2. MRI markers of glymphatic dysfunction are largely confined to an inflammatory subgroup of schizophrenia

Substantial evidence from human imaging and animal models indicates that both peripheral and central inflammation can impair glymphatic clearance [29–32]. There is also converging evidence for an inflammatory subtype of SCZ that can be identified by elevated peripheral inflammatory markers [14, 33] or neuroinflammatory signatures in post-mortem brain tissue [15–20]. However, it remains unclear whether glymphatic dysfunction in SCZ is primarily driven by inflammation.

To address this issue, we examined the associations between peripheral inflammatory markers and DTI-ALPS index / CPV in our patient cohort. Inflammatory markers assessed included CRP, IL-6, total leukocytes, percentages of neutrophils, monocytes, lymphocytes, and platelets, as well as derived cellular ratios, including the neutrophil-to-lymphocyte ratio (NLR), monocyte-to-lymphocyte ratio (MLR), platelet-to-lymphocyte ratio (PLR), systemic immune-inflammation index (SII: platelets × neutrophils/lymphocytes), systemic inflammation response index (SIRI: neutrophils × monocytes/lymphocytes), and pan-immune-inflammation value (PIV: platelets × neutrophils × monocytes/lymphocytes). Since we expected strong correlations among inflammatory markers, we first conducted a principal component analysis (PCA) to reduce the burden of multiple testing (Figure 2B). PCA revealed three principal components that captured ∼ 82% of variance. The first principal component was defined by the relationship between monocytes, neutrophils, and platelets on the one hand and lymphocytes on the other hand representing a proxy of the relationship between innate and adaptive immunity. This component was negatively correlated with DTI-ALPS index (p-FDR < 0.05) and positively correlated with CPV, although the latter finding showed only a trend-level when correcting for multiple comparisons (p-FDR = 0.084). The second principal component was largely defined by monocytes and did not show any significant correlation with DTI-ALPS index or CPV. The third principal component was characterized by CRP and IL-6 levels and exhibited a significant negative correlation with DTI-ALPS index (p-FDR < 0.05), whereas the association with CPV reached trend-level significance only (p-FDR = 0.121). Because we expected a reduced DTI-ALPS index and increased CPV to be confined to patients with higher inflammatory marker levels, we conducted subgroup analyses in patients with SCZ stratified by the median of the first principal component (Figure 2D) and the third principal component (Figure 2E). For both principal components, DTI-ALPS index decreased, whereas CPV increased, in a stepwise manner from healthy controls to patients in the low-inflammatory group and further to those in the high-inflammatory group, indicating a graded, inflammation-related effect. Moreover, significant differences relative to healthy controls were observed only in patients in the high-inflammatory subgroups. Taken together, imaging measures of glymphatic dysfunction in our SCZ cohort were predominantly found in patients with elevated peripheral inflammatory markers, particularly those reflecting activation of the innate immune system.

**Figure 2.**
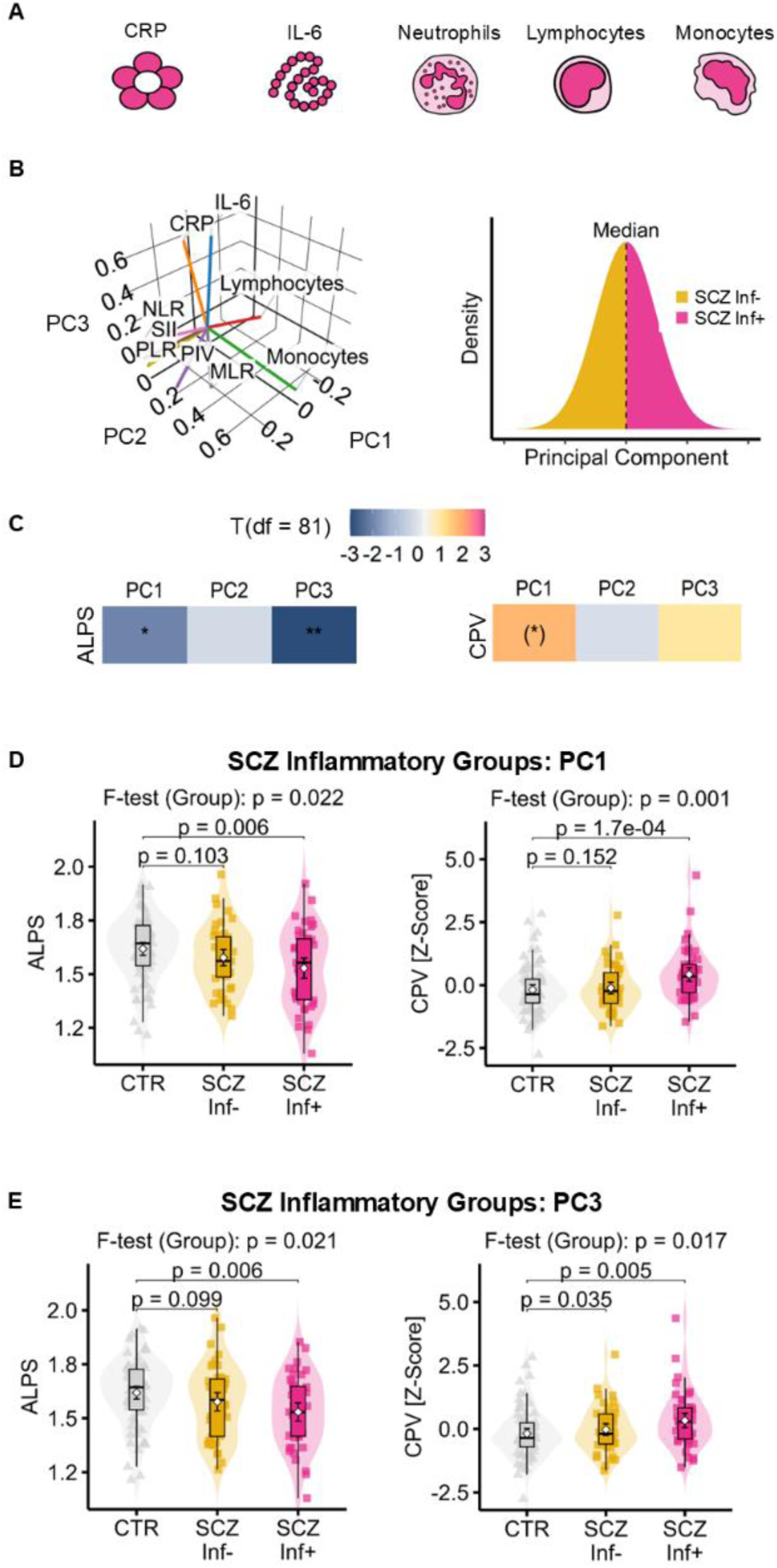
A.Illustration of the different peripheral inflammatory markers assessed in patients with schizophrenia. Platelets are not shown for simplification. Further inflammatory markers included multiple derived cellular ratios (see main manuscript). B.Given the anticipated strong intercorrelations among the inflammatory markers, we first applied principal component analysis (PCA) to limit the burden of multiple testing. Left panel: The first three principal components are shown as orthogonal axes in 3D space, with inflammatory marker loadings visualized as vectors. For clarity, only inflammatory markers with a loading of at least 0.336 on at least one of the three principal components are displayed. The first principal component was characterized by the relationship between monocytes, neutrophils, and platelets on the one hand and lymphocytes on the other. Accordingly, the derived ratios (NLR, MLR, PLR, PIV, SIRI, SII) and, to a lesser extent, total leukocytes, monocytes, neutrophils, and platelets loaded positively on this component, whereas lymphocytes showed a negative loading. To assess the relationship between the inflammation-related principal components and the DTI-ALPS index and CPV, we conducted separate analyses treating each principal component either as a continuous variable (C) or as a categorical variable with two levels (low vs high inflammation) defined by a median split (right panel of B, D-E). The latter approach enabled us to compare each subgroup to the control subjects. C. Linear models were fitted with the DTI-ALPS index or CPV as the dependent variable and the respective PC as the independent (continuous) variable of interest. Age, gender, total intracranial volume, and antipsychotic burden were entered as independent variables of no interest. For CPV, robust linear models were used because of the presence of outliers. One-sided p values were employed, as we hypothesized that higher levels of inflammatory markers would be associated with a lower DTI-ALPS index and higher CPV. *: p-FDR < 0.05; **: p-FDR < 0.01; (*): p < 0.05, p-FDR > 0.05 D. -E. Inflammatory subgroups of SCZ were defined based on a median split of PC1 (D) or PC3 (E). DTI-ALPS index (left panel) or CPV (right panel) were compared between each subgroup of patients and the control subjects using linear models (DTI-ALPS index) or robust linear models (CPV) including age, gender, and total intracranial volume as covariates of no interest. Robust linear models were chosen for CPV because of the presence of outliers. Overall group effects were assessed using F-tests for linear models and F-test-like Wald’s tests for robust linear models. Post hoc group comparisons were conducted using Dunnett’s tests for linear models and estimated marginal means contrasts with false discovery rate (FDR) correction of p-values for robust linear models. For both DTI-ALPS index and CPV, a gradual decrease / increase was observed across healthy subjects, low and high inflammation groups. Statistically significant decreases in the DTI-ALPS index relative to the healthy control group were observed only in the high-inflammation groups, both under the PC1-based (D) and PC3-based (E) group definitions. In a similar manner, under the PC1-based group definition, a statistically significant increase in CPV relative to the healthy control group was observed only in the high-inflammation group (D). Under the PC3-based group definition, both the low- and high-inflammation subgroups exhibited significantly higher CPV values than the healthy control group (E).

### 2.3 MRI markers of glymphatic dysfunction are associated with schizophrenia polygenic risk enriched for immune system and oxidative stress-related aging pathways

Schizophrenia is a highly polygenic disorder including common variants with genome wide significance at 287 loci according to the most recent meta-analytic evidence [34]. Using Gene Ontology (GO) annotations, we assigned all genes mapped to these 287 loci to two biological pathways relevant to our mechanistic hypothesis of glymphatic dysfunction, namely immune-related processes and aging. Loci whose mapped genes could not be assigned to either pathway were grouped into a remainder (“rest”) category (Figure 3 A & B).

**Figure 3.**
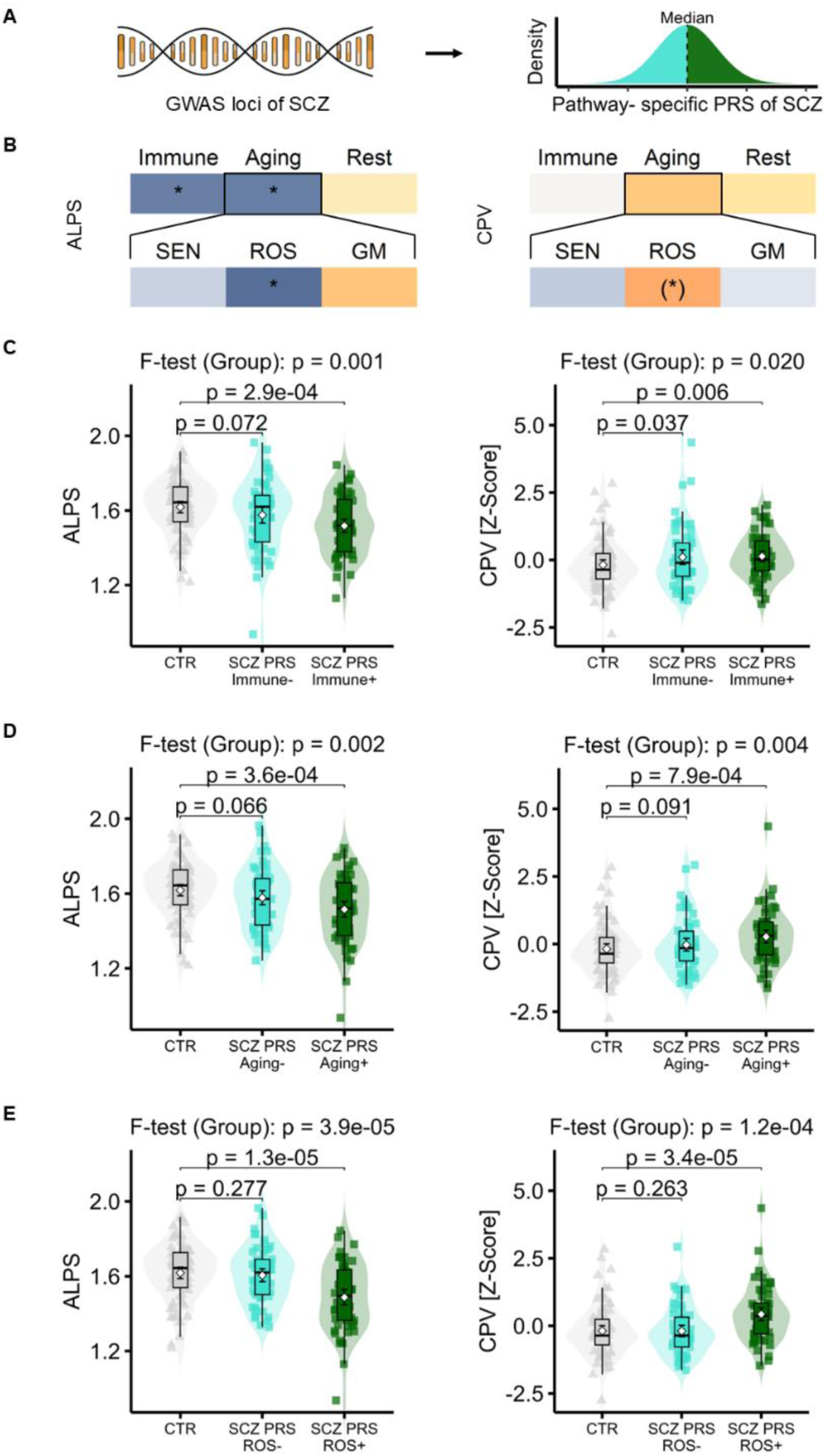
A. Pathway-specific polygenic risk scores (PRS) of SCZ were defined by mapping genes associated with the 287 genome-wide significant loci from the latest GWAS of SCZ to immune-related processes and aging, using GO terms. Loci whose mapped genes did not match either pathway were grouped into a remainder (“rest”) category. To assess the relationship between pathway-specific PRS scores and the DTI-ALPS index and CPV, we conducted separate analyses treating pathway-specific PRS scores either as a continuous variable (see panel B) or as a categorical variable with two levels (low vs high PRS burden) defined by a median split (right panel of A, C-E). B. Treating pathway-specific PRS either as a continuous variable, we fitted linear models with the DTI-ALPS index or CPV as the dependent variable and the respective pathway-specific PRS as the independent variable of interest. Age, gender, total intracranial volume, and antipsychotic burden were entered as independent variables of no interest. For CPV, robust linear models were used because of the presence of outliers. Since aging-related PRS was defined by GO terms related to senescence (SEN), reactive oxygen species (ROS) and genome maintenance (GM), we also did separate analyses for each of these sub-pathways, too. One-sided p values were employed, as we hypothesized that higher levels of pathway-specific PRS scores would be associated with a lower DTI-ALPS index and higher CPV. *: p-FDR < 0.05; (*): p < 0.05, p-FDR > 0.05 C. - E. Subgroups of SCZ were defined based on a median-split of the immune-(C), aging-(D) or ROS-related (E) PRS of SCZ. DTI-ALPS index (left panel) or CPV (right panel) were compared between each subgroup of patients and the control subjects using linear models (DTI-ALPS index) or robust linear models (CPV) including age, gender, and total intracranial volume as covariates of no interest. Robust linear models were chosen for CPV because of the presence of outliers. Overall group effects were assessed using F-tests for linear models and F-test-like Wald’s tests for robust linear models. Post hoc group comparisons were conducted using Dunnett’s tests for linear models and estimated marginal means contrasts with false discovery rate (FDR) correction of p-values for robust linear models. For both DTI-ALPS index and CPV, a gradual decrease / increase was observed across healthy subjects, low and high PRS groups. Statistically significant decreases in the DTI-ALPS index relative to the healthy control group were observed only in the high-PRS groups, including the immune- (C), aging-(D) and ROS-based (E) group definitions. In a similar manner, both under the aging- and ROS-based group definition, statistically significant increases in CPV relative to the healthy control group were observed only in the high-PRS groups (D & E). Under the immune-based group definition, both the low- and high-PRS subgroups exhibited significantly higher CPV values than the healthy control group (C).

Indeed, lower DTI-ALPS index was significantly correlated with higher SCZ polygenic risk scores related to the immune system (PRS-immune) and aging (PRS-aging), but not the remaining polygenic risk of SCZ as represented by the remainder category (Figure 3B, left panel). Since PRS-aging was constructed by different GO terms, i.e. “senescence”, “genome maintenance” and “reactive oxygen species” (ROS), we also assessed polygenic risk scores related to these sub-categories. Here, lower DTI-ALPS index was significantly correlated with higher ROS-related PRS. CPV, in contrast, did not show any significant correlation with any of the three main PRS-categories (Figure 3B, right panel). For the subcategories of PRS-aging, CPV exhibited a positive association with PRS-ROS, which showed a trend-level significance after FDR-correction, only (p-FDR = 0.078).

As a complementary analysis, we also treated each pathway-related PRS that showed any significant correlation with the DTI-ALPS index or CPV as a categorical variable with two levels (low vs high PRS burden) defined by a median split and compared each subgroup to the healthy control group (Figure 3 C – E). Similar to the findings in 2.2, we observed a stepwise decrease in DTI-ALPS index and a stepwise increase in CPV from healthy controls to patients in the low-PRS group, and further to those in the high-PRS group, under the immune-, aging-, and ROS-based group definitions. Statistically significant reductions in the DTI-ALPS index relative to healthy controls emerged only in the high-PRS groups, across the immune-(Figure 3 C), aging-(Figure 3 D), and ROS-based (Figure 3 E) group definitions. Similarly, CPV was significantly increased relative to healthy controls only in the high-PRS groups under the aging- and ROS-based definitions (Figure 3 D and E), whereas under the immune-based definition both the low-and high-PRS subgroups showed significantly higher CPV values than healthy controls (Figure 3 C).

### 2.4 Structural equation modeling indicates that glymphatic dysfunction driven by inflammation contributes to cognitive impairment in schizophrenia

Because our mechanistic framework postulated that glymphatic dysfunction mediates the association between inflammation and cognitive impairment in schizophrenia, we employed structural equation modeling (SEM). Since we assumed that choroid plexus enlargement reflects hypertrophy and structural remodeling as a direct consequence of impaired glymphatic CSF flow, our primary model postulated that impaired perivascular CSF flow (approximated by DTI-ALPS index) precedes increased CPV which in turn leads to cognitive impairment (quantified by the total score of the Brief Assessment of Cognition in Schizophrenia, BACS). Peripheral inflammation (quantified by the z-standardized mean of the z-standardized first and third principal component of the assessed peripheral inflammatory markers) was considered to influence both perivascular CSF flow and CPV. Our primary model showed excellent fit (Fig. 4, Supplementary Figure 1, first row, right panel), whereas models assuming that CPV precedes reduced DTI-ALPS index showed clearly inferior fits (Supplementary Figure 1, second row). Removing a path from inflammation to CPV in our primary model did not significantly worsen model fit (Supplementary Figure 1, first row, right panel), whereas removing a path from inflammation to the DTI-ALPS index in the reversed-order model (CPV preceding impaired perivascular CSF flow) significantly worsened it, underscoring the importance of this pathway (Supplementary Figure 1, second row, right panel). Accordingly, the simplest model to describe the data suggests that inflammation drives impaired perivascular CSF flow which subsequently mediates choroid plexus enlargement that finally leads to cognitive impairment.

**Figure 4.**
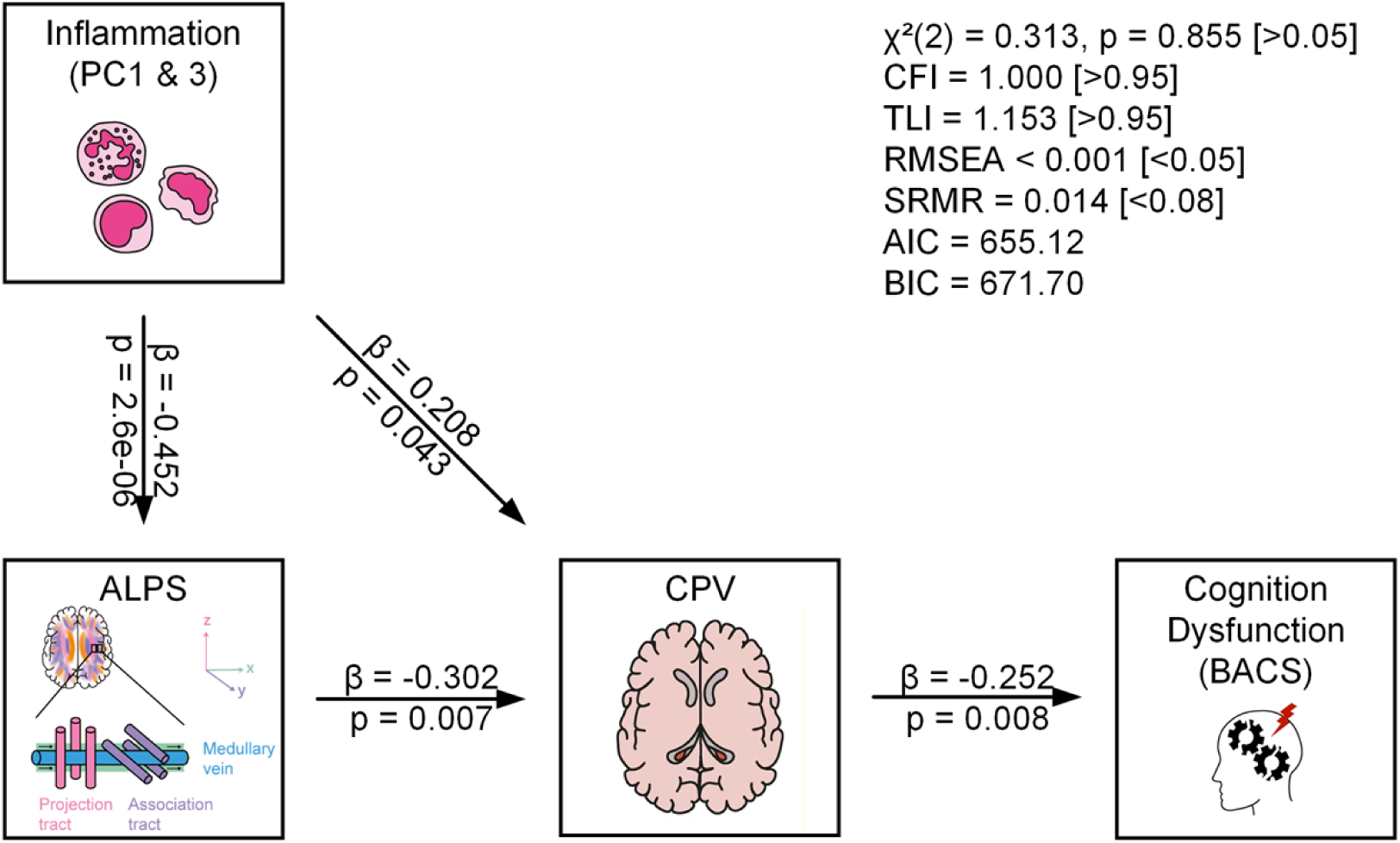
Structural equation modeling. Our structural model included four variables: peripheral inflammatory burden (mean of the z-standardized scores of PC1 and PC3), the DTI-ALPS index, CPV, and the BACS composite score, arranged in a hypothesized serial pathway from inflammation to cognitive impairment. P-values related to path coefficients are one-sided, as we hypothesized that higher levels of inflammatory markers would be associated with a lower DTI-ALPS index and higher CPV, and that higher CPV would be associated with poorer cognitive performance. Fit indices reported in the upper right corner suggest an excellent model fit. CFI — Comparative Fit Index, TLI — Tucker–Lewis Index (also called the Non-Normed Fit Index, NNFI), RMSEA — Root Mean Square Error of Approximation, SRMR — Standardized Root Mean Square Residual, AIC — Akaike Information Criterion, BIC — Bayesian Information Criterion

### 2.5. Unbiased clustering of brain transcriptomic data identifies a neuro-inflammatory subgroup enriched in SCZ

To better understand the transcriptomic basis of inflammation and to identify potential inflammatory subgroups of individuals with SCZ, we analyzed post-mortem dorsolateral prefrontal cortex (DLPFC) and orbitofrontal cortex (OFC) transcriptomic data from two published datasets [32, 35], comprising a total of 92 control donors and 95 donors with SCZ (Figure 5A). As a first step, we applied an unbiased clustering procedure to enable the identification of individuals with neuroinflammation in patients with SCZ and control subjects. Joint clustering of microglial and astrocytic pseudo-bulk expression profiles revealed a clear separation of two major clusters consisting of samples with high inflammation (Inf+) and low inflammation (Inf-), characterized by the inverse expression of inflammatory and homeostatic markers in both microglia and astrocytes (Figure 5B, C, Supplementary Figure 2A).

**Figure 5.**
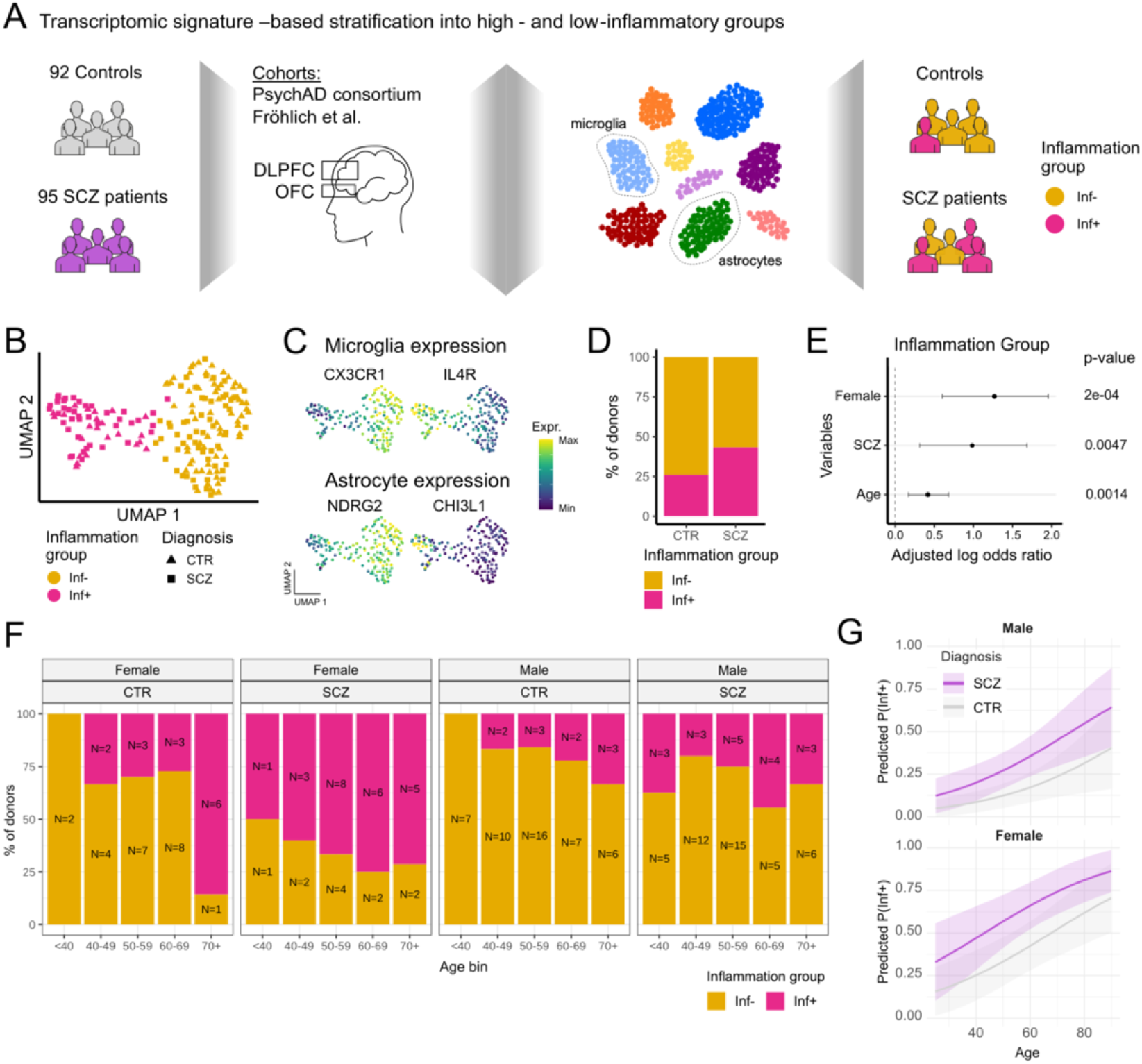
High inflammation profile defined by microglial and astrocyte states is enriched in schizophrenia. A. Schematic overview of the post-mortem study design. Pseudobulk transcriptomic profiles of microglia and astrocytes were derived from published post-mortem dorsolateral prefrontal cortex (DLPFC) and orbitofrontal cortex (OFC) datasets and used for the unbiased stratification of donors into low-inflammatory (Inf−) and high-inflammatory (Inf+) subgroups. B. Joint UMAP embedding of microglial and astrocytic pseudobulk transcriptomes coloured by inflammatory group (Inf−, yellow; Inf+, pink). Point shape denotes diagnosis (controls (CTR), triangles; schizophrenia (SCZ), squares). C. Feature plots showing normalized expression levels of representative homeostatic and inflammatory markers in microglia (CX3CR1 and IL4R) and astrocytes (NDRG2 and CHI3L1). D. Distribution of CTR and SCZ donors across inflammatory subgroups. Bar heights indicate the percentage of donors assigned to each subgroup. E. Multivariable logistic regression analysis of factors associated with assignment to the Inf+ subgroup. Diagnosis, gender and age were included as explanatory variables in the model. Points indicate adjusted log-odds ratios, and lines represent 95% confidence intervals. Coefficient P values are indicated. F. Distribution of donors assigned to the Inf− and Inf+ subgroups, stratified by diagnosis, sex, and binned age group. Bar heights represent percentages, and numbers within bars denote donor counts. G. Predicted probability of assignment to the Inf+ subgroup according to diagnosis, sex, and age derived by multivariable logistic regression. Lines represent model predictions and shaded areas indicate 95% confidence intervals.

43% of SCZ patients were assigned to the Inf+ group, while only 26% of control donors were assigned to this group (Figure 5D), indicating a marked enrichment of neuroinflammatory signatures among individuals with SCZ. To identify factors associated with the Inf+ phenotype, we then applied multivariate logistic regression to assess the association of technical and clinical attributes with the Inf+ group. This analysis revealed that Inf+ samples were associated with older age and a higher proportion of female donors, while source cohort, post-mortem interval, and tissue pH showed no significant effects (Figure 5E-F, Supplementary Figure 2B). Furthermore, our model predicted a consistently higher probability of having an Inf+ phenotype in both male and female SCZ patients compared to controls across all ages (Figure 5G). Patients with SCZ in general exhibited probabilities of inflammatory phenotypes that were found in the controls only 20-30 years later. This pronounced age shift is consistent with the concept of “inflammaging” and suggests that individuals with SCZ may exhibit an accelerated age-related neuroinflammatory trajectory [36].

### 2.6. A transcriptomic brain signature of glymphatic dysfunction is detected in a high inflammatory subgroup of SCZ

At the molecular level, glymphatic brain clearance relies on the polarized localization of AQP4 channels at astrocytic endfeet, which is mediated by the dystrophin-associated protein complex (DAPC) that anchors AQP4 to the perivascular basal lamina (Fig. 6A). Disruption of this complex has been shown to dramatically impair CSF–interstitial fluid exchange and waste clearance [37]. However, expression changes in DAPC components have not yet been studied in post-mortem samples of patients with SCZ. Based on our findings in the in vivo cohort, we hypothesized a strong dysregulation of genes belonging to the dystrophin-associated complex in SCZ – particularly in patients with inflammatory phenotype (Inf+ patients).

**Figure 6.**
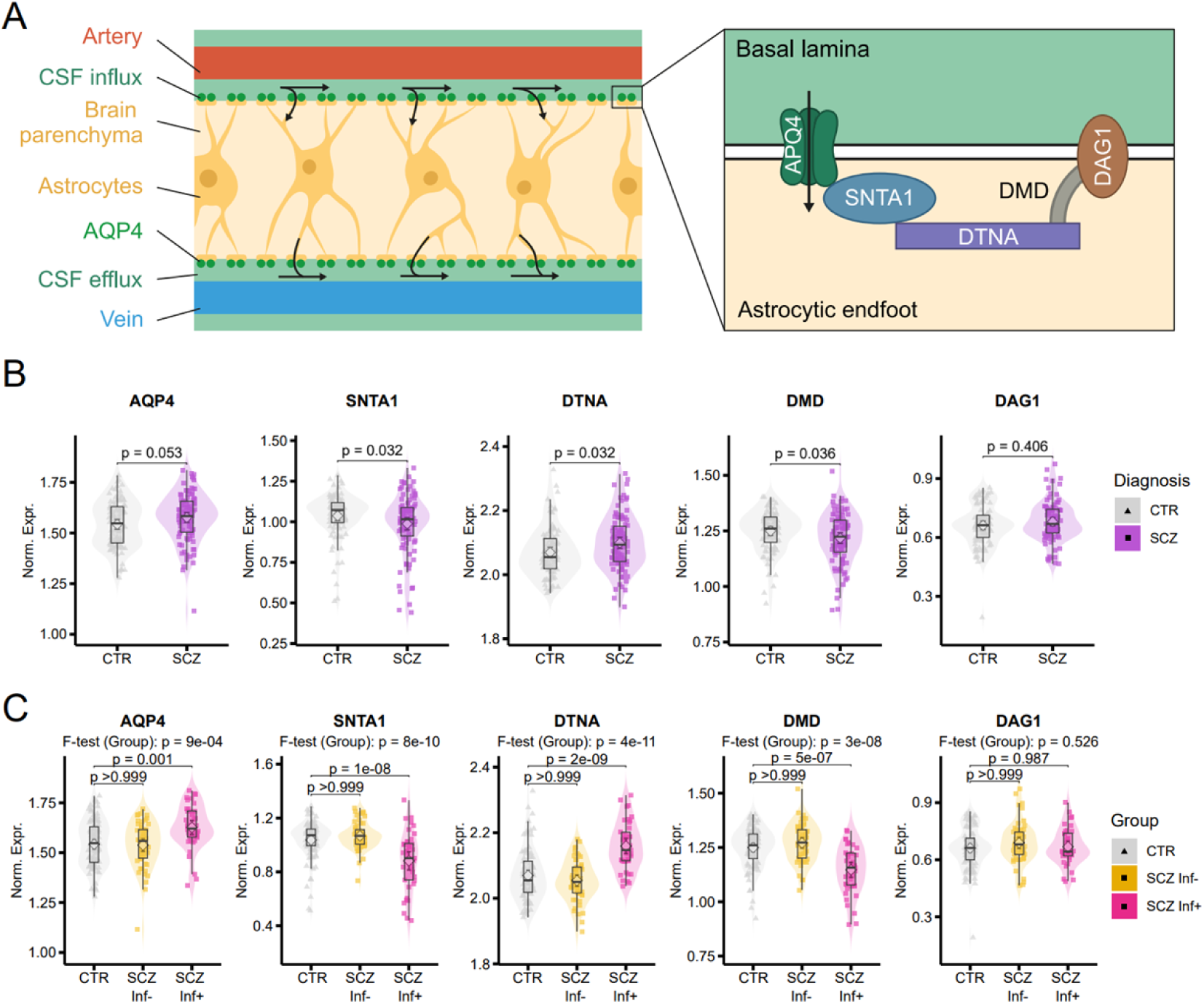
Altered expression of glymphatic function-associated genes is confined to the high-inflammatory subgroup of schizophrenia. **A.** Schematic illustration of the astrocytic endfoot–basal lamina interface, highlighting key components of the dystrophin-associated protein complex (DAPC), including DAG1, DMD, DTNA, and SNTA1 as well as the polarized localization of the water channel AQP4,. **B.** Normalized astrocytic expression of AQP4, SNTA1, DTNA, DMD, and DAG1 in control (CTR, grey) and schizophrenia (SCZ, purple) donors. Individual points represent pseudobulk expression values. Violin plots depict expression distributions and boxplots indicate median and interquartile range. Central points and error bars represent group means ± 95% confidence intervals. Statistical significance was assessed using type II ANOVA F-tests with diagnosis as the explanatory variable. Displayed P values are two-sided and false discovery rate (FDR)-corrected. **C.** Normalized astrocytic expression of AQP4, SNTA1, DTNA, DMD, and DAG1 stratified by inflammatory subgroup. Individual points represent pseudobulk expression values. Violin plots depict expression distributions and boxplots indicate median and interquartile range. Central points and error bars represent group means ± 95% confidence intervals. Overall group effects were assessed using F-tests; displayed P values are two-sided and FDR-corrected. Post hoc comparisons were performed using Dunnett’s tests with CTR as the reference group.

To test this hypothesis, we examined astrocytic expression levels of AQP4 and the DAPC (DAG1, DMD, DTNA, SNTA1) comparing the full SCZ cohort as well as low- and high-inflammation subgroups with healthy controls (Figure 6A). Across all patients with SCZ, expression of both DMD and SNTA1 was significantly reduced (Figure 6B), the latter of which is a direct binding partner of AQP4 at astrocytic endfeet [37, 38]. Moreover, a significant increase of DTNA-expression and a tendential increase of total AQP4 expression (p-FDR = 0.053) were detected, whereas DAG1 expression was not altered. Strikingly, when SCZ samples were separated by inflammatory status, this pattern became much more pronounced in the Inf+ group, whereas Inf-SCZ samples were largely indistinguishable from controls (Figure 6C). This pattern suggests impaired stabilization of perivascular AQP4 under conditions of elevated inflammation.

For further downstream analyses, we integrated the dysregulated genes related to AQP4 and the DAPC in SCZ into a single glymphatic dysregulation score (Figure 7A).

**Figure 7.**
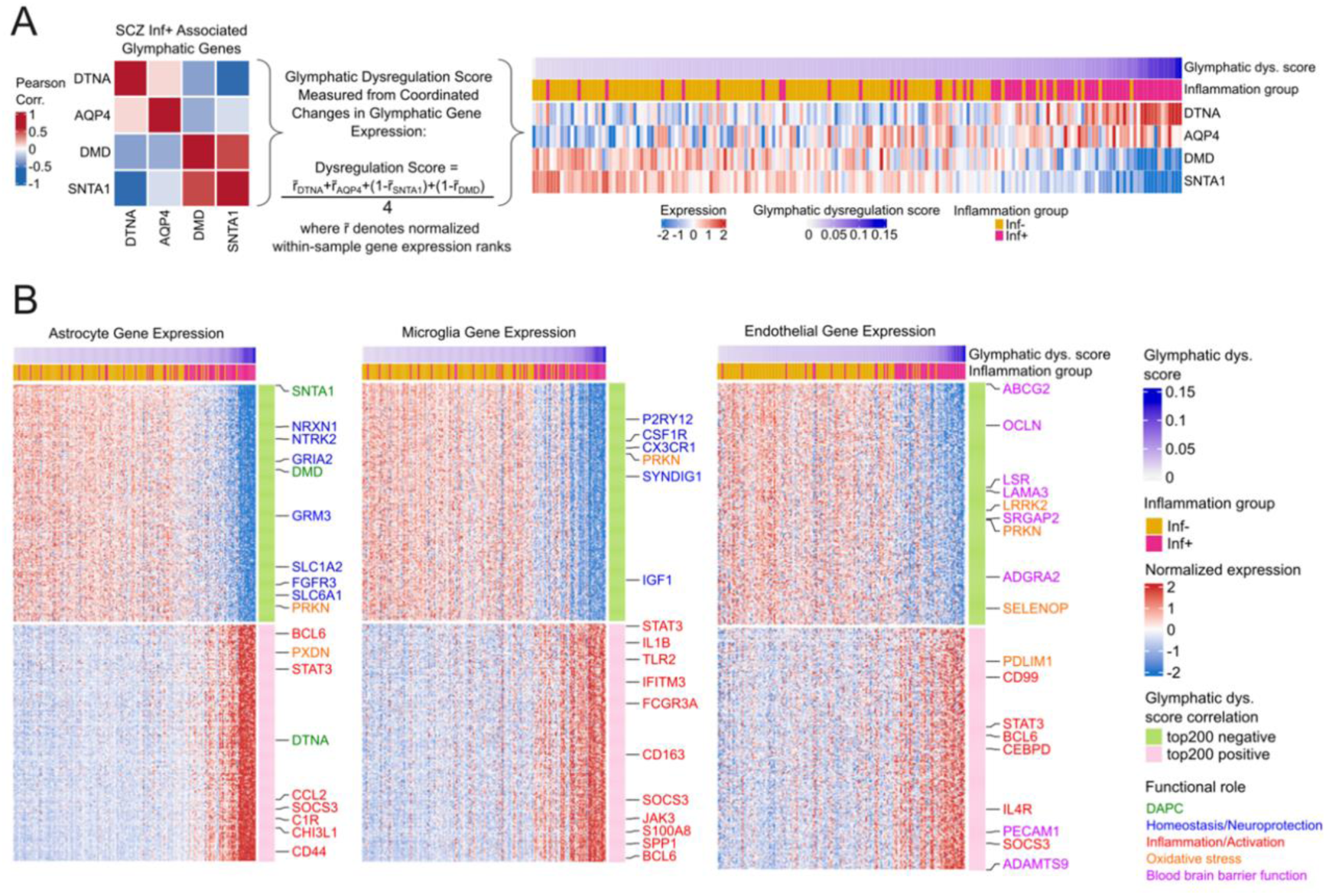
A glymphatic dysregulation score reveals coordinated molecular alterations associated with the high-inflammatory schizophrenia subgroup. **A.** Construction of a glymphatic dysregulation score based on expression patterns of glymphatic function-associated genes altered in SCZ. The left heatmap shows Pearson correlation coefficients between astrocytic expression of DTNA, AQP4, DMD, and SNTA1. A glymphatic dysregulation score was calculated from within-sample normalized expression ranks of these four genes, capturing coordinated transcriptional changes related to astrocytic endfoot organization as a proxy of glymphatic function. The right heatmap shows normalized gene expression across donors ordered by increasing glymphatic dysregulation score. Annotation bars indicate glymphatic dysregulation score and inflammatory subgroup assignment (Inf−, yellow; Inf+, pink). Expression values are displayed as row-wise z-scores. **B.** Cell type-specific transcriptional correlates of glymphatic dysregulation. Heatmaps show normalized expression of the 200 genes most negatively correlated (green) and the 200 genes most positively correlated (pink) with the glymphatic dysregulation score in astrocytes, microglia, and endothelial cells. Donors are ordered by increasing glymphatic dysregulation score. Selected representative genes are indicated and colored by functional role. Annotation bars denote glymphatic dysregulation score and inflammatory subgroup assignment. Expression values are displayed as row-wise z-scores.

Next, we identified genes beyond the selected marker genes whose expression was correlated or anticorrelated with the glymphatic signature score, not only within astrocytes, but also within microglia and endothelial cells (Figure 7B, Supplementary Table 2). Genes most strongly anti-correlating with the glymphatic dysregulation score were enriched for homeostatic and neuro-supportive function in astrocytes (NRXN1, NTRK2, FGFR3) [39–41] and microglia (P2RY12, CX3CR1, CSF1R) [39, 42, 43]. Furthermore, several genes involved in oxidative stress and mitochondria function (PRKN, LRRK2, SELENOP, PDLIM1, PXDN) [44–48] were dysregulated across multiple cell types, in line with our previous observation of higher ROS-related polygenic risk scores in SCZ patients with decreased DTI-ALPS and increased CPV (Figure 3B, E). In contrast, genes involved in immune activation and chronic inflammation -including canonical NF-κB target genes - (CCL2, BCL6, JAK3, STAT3, IL1B) [49, 50] were strongly correlated with the glymphatic dysregulation score across multiple cell types. Notably, endothelial cells displayed transcriptional hallmarks of blood-brain barrier dysfunction including down-regulation of tight junction proteins (OCLN, LSR, SRGAP2, ADGRA2) [51, 52], in conjunction with elevated inflammation and leukocyte migration markers (PECAM1, CD99, STAT3, BCL6) [53], indicating a reduction in blood-brain barrier integrity in SCZ patients with higher glymphatic dysfunction. Together, these data indicate that glymphatic dysfunction emerges in specific cell-associated transcriptional programs in the context of heightened inflammation.

### 2.7. Multimodal evidence for glymphatic dysfunction as a mediator of accelerated aging in schizophrenia

To further study the relationship between impaired glymphatic brain clearance and accelerated aging in SCZ, we assessed aging-related gene expression signatures in the post-mortem datasets as well as brain age and DNA methylation age gaps in the in vivo dataset (Figure 8A). In the post-mortem brain tissue, we studied aging signatures for microglia and astrocytes using genes whose expression was shown to be associated with donor age [54]. Indeed, for both astrocytes and microglia, the aging signature score showed a significant positive linear relationship with donor age (Figure 8B, first and third panel). SCZ patients with above median glymphatic dysregulation score (SCZ glymph-) had consistently higher aging signature scores compared to controls and SCZ glymph+ patients in both cell types (Figure 8B, second panel and fourth panel), suggesting accelerated aging in patients with glymphatic impairment.

**Figure 8.**
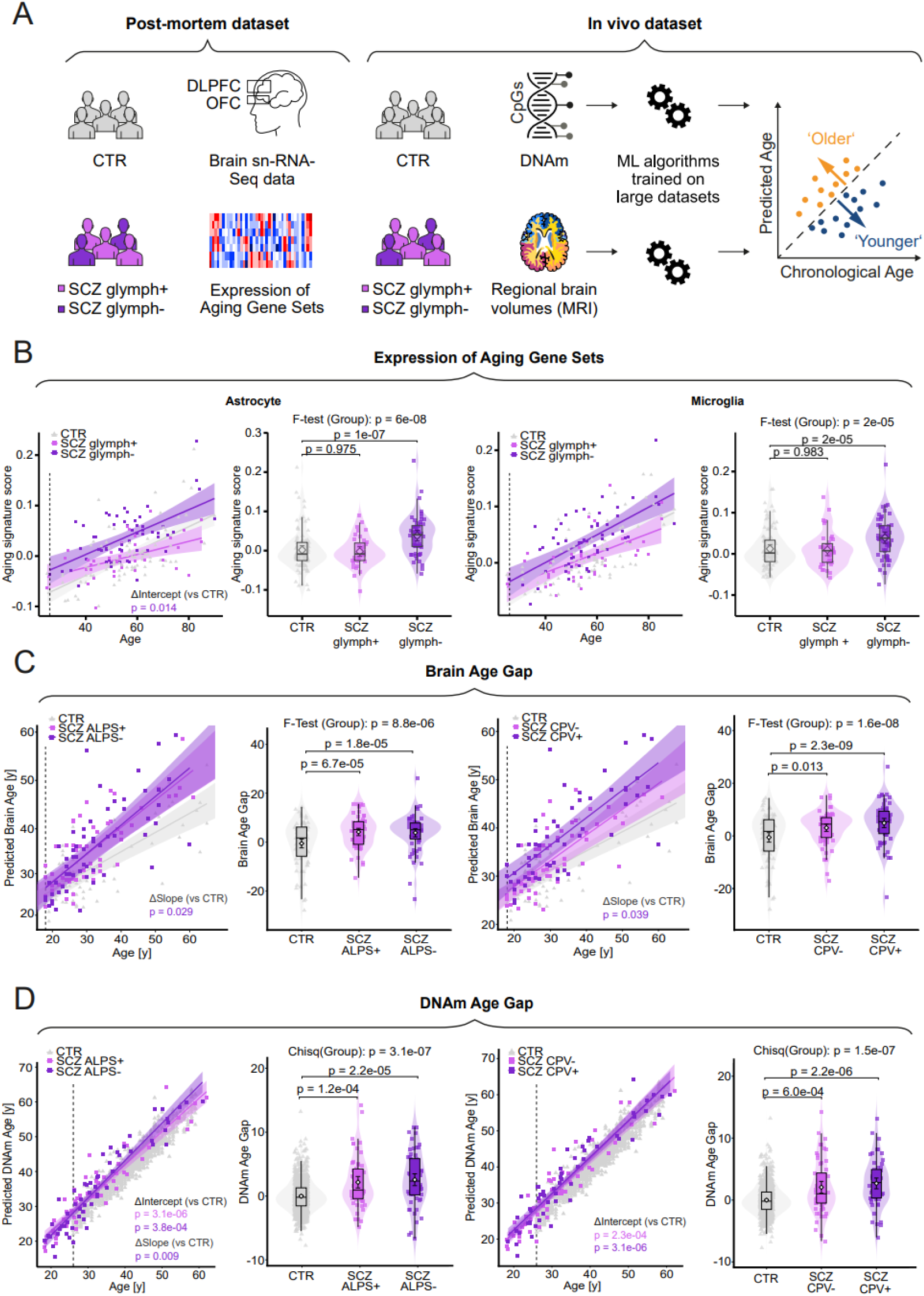
Convergent transcriptomic, neuroimaging, and epigenetic evidence for accelerated aging in schizophrenia subgroups defined by glymphatic dysfunction. A. Aging-related signatures were assessed across complementary post-mortem and in vivo datasets. In post-mortem dorsolateral prefrontal cortex (DLPFC) and orbitofrontal cortex (OFC) single-nucleus RNA-sequencing datasets, aging signature scores were derived from previously established age-associated gene sets in astrocytes and microglia. Patients with schizophrenia (SCZ) were stratified according to glymphatic dysfunction status based on the glymphatic dysregulation score described in Fig. 6. In the in vivo cohort, biological aging was quantified using DNA methylation (DNAm) age and MRI-derived brain age prediction models. Accelerated aging was evaluated as the deviation between predicted biological age and chronological age (age gap). B-D. Solid lines in scatter plots indicate fitted linear regression models and shaded areas denote 95% confidence intervals. Adjacent violin plots display group distributions; boxplots indicate median and interquartile range (IQR), and points with error bars denote group means ± 95% CI for two-tailed and ± 90% CI for one-tailed p-values. Statistical tests and post hoc comparisons (Dunnett’s, CTR reference) are indicated in each panel. B. Aging-related transcriptomic signatures in astrocytes and microglia across controls (CTR), SCZ glymph+, and SCZ glymph− donors. Scatter plots show aging signature scores as a function of donor age. For both astrocytes and microglia, SCZ glymph− donors show significantly higher aging signatures than controls, whereas SCZ glymph+ donors are largely indistinguishable from controls. Group effects were assessed by F-tests followed by Dunnet’s test to compare each SCZ subgroup with the CTR group. Indicated p-values are two-tailed. C. MRI-derived brain age predictions and brain age gaps in controls (CTR), SCZ ALPS+, and SCZ ALPS− (left), and controls, SCZ CPV−, and SCZ CPV+ (right). Group effects were assessed by F-tests followed by Dunnet’s tests to compare each SCZ subgroup with the CTR group. Indicated p-values of Dunnet’s tests are right-tailed as we expected higher brain age gaps in patients with SCZ based on the literature. D. DNA methylation (DNAm) age predictions and DNAm age gaps in controls (CTR), SCZ ALPS+, and SCZ ALPS− (left), and controls, SCZ CPV−, and SCZ CPV+ (right). Group effects were tested using Wald χ² statistics from linear models with HC3 robust standard errors to account for unequal variances. Estimated marginal means contrasts with FDR-correction were applied to compare each SCZ subgroup with the CTR group. Indicated p-values of Dunnet’s tests are right-tailed as we expected higher brain age gaps in patients with SCZ based on the literature.

Growing evidence suggests that accelerated brain aging in SCZ can be studied using brain volume changes as quantified by MRI [5]. To explore whether MRI-based brain age gaps are related to impaired glymphatic brain clearance, we compared brain age gap between healthy controls and patients with SCZ who were stratified by DTI-ALPS index or CPV. Individual brain age estimates were obtained using a cross-validated support vector regression (SVR) model trained on regional grey matter and CSF volumes from healthy controls (n = 73), including an external cohort with a broad age range (n = 3227) [55]. Brain age gap was calculated after correcting for age-related bias using linear regression and SCZ subgroups were defined based on a median split of DTI-ALPS index or CPV. As expected, a positive linear relationship between chronological age and the bias-corrected predicted brain age emerged for each group – both for the DTI-ALPS (Figure 8C – first panel) or CPV-based group definition (Figure 8C – third panel). A stepwise increase in the brain age gap was observed from controls to the high-ALPS/low-CPV group and further to the low-ALPS/high-CPV group, with both SCZ subgroups differing significantly from healthy controls (Figure 8C, second and fourth panel). However, only the low-ALPS/high-CPV group showed a significantly steeper increase in brain age over time (i.e., a higher slope of bias-corrected brain age versus chronological age), consistent with accelerated brain aging in SCZ patients with glymphatic dysfunction (Figure 8C, second and fourth panel).

Besides evidence for accelerated brain aging, a growing number of studies have also revealed elevated DNA methylation age in patients with SCZ [9, 10]. Again, the relationship with glymphatic clearance has not been addressed yet. We generated DNA methylation data from patients’ whole blood using Illumina EPIC arrays, applied standard quality control and preprocessing, and computed the first principal component of GrimAge, a machine-learning–derived epigenetic clock trained on time-to-death that surpasses earlier clocks in forecasting both disease risk and mortality [56]. Since no blood samples were available for our healthy control group, we utilized an external reference data set comprising 1024 healthy subjects [57]. Similar to the brain age gap, the DNAm age gap was calculated after correction for age-related bias using linear regression, and SCZ subgroups were defined based on a median split of the DTI-ALPS index or CPV. In all groups, DNAm age showed a positive linear relationship with chronological age (Figure 8D, first and third panels).

When stratified by DTI-ALPS index, both SCZ subgroups exhibited significantly increased DNAm age gaps compared to healthy controls, with the largest effect observed in the DTI-ALPS− group (below median). Moreover, only the DTI-ALPS− group showed a significantly steeper increase in DNAm age over time (i.e., a higher slope of DNAm age versus chronological age) compared to controls, whereas the DTI-ALPS+ group did not. Stratification by CPV revealed significantly elevated DNAm age gaps in both SCZ subgroups relative to controls, with the highest values observed in the CPV+ group. Both subgroups showed significantly higher intercepts of the DNAm age versus chronological age regression compared to controls, while the slopes were not significantly different. Taken together, these findings provide convergent evidence across datasets and modalities for accelerated aging trajectories in SCZ patients with signatures of glymphatic dysfunction.

## 3. Discussion

Reduced life expectancy and persistent cognitive impairment remain among the most pressing unresolved challenges in SCZ. Our findings implicate dysfunction of the glymphatic system—the brain’s waste clearance system—as a central mechanism linking neuroinflammation to accelerated biological aging and cognitive decline. Glymphatic clearance relies on the polarized expression of AQP4 channels at astrocytic endfeet which is mediated by the DAPC that anchors AQP4 to the perivascular basal lamina. As a major finding of the present study, we identify a dysregulated transcriptomic signature involving several components of the AQP4-DAPC complex in post-mortem brain samples from patients belonging to an inflammatory subgroup of SCZ. This signature was characterized by a downregulation of AQP4’s direct binding partner SNTA1 as well as DMD, and upregulation of DTNA and AQP4. Consistent with this pattern, knockout of different components of the DAPC (e.g. SNTA1) in mice has been shown to lead to substantially reduced AQP4 expression at astrocytic endfeet and impaired glymphatic clearance while increasing AQP4 in other membrane domains [37, 38].

Mechanistic animal and associative human studies increasingly link impaired glymphatic clearance to inflammation, [29–31, 58, 59], yet this relationship has not been systematically investigated in patients with SCZ, despite substantial evidence implicating immunological mechanisms in its pathophysiology. Immune-related risk factors and inflammatory mechanisms in SCZ span prenatal or early-life infection and maternal immune activation, as well as dysregulated microglia-mediated synaptic pruning during adolescence and early adulthood [60, 61]. Moreover, one of the most robust and consistently replicated schizophrenia GWAS signals maps to the extended major histocompatibility complex (MHC) region, further supporting the involvement of immune-related pathways in disease susceptibility [34, 62]. Peripheral inflammatory profiling and post-mortem brain transcriptomics independently identify subgroups of patients with SCZ characterized by heightened inflammatory activity [16, 19, 20, 63–66]. In the present study, we show that – across modalities - these independently defined high-inflammatory subgroups of SCZ converge on signatures of glymphatic dysfunction and accelerated biological aging. Peripheral inflammatory markers that were most strongly associated with imaging indices of glymphatic dysfunction included elevated neutrophil-to-lymphocyte and monocyte-to-lymphocyte ratios, as well as increased IL-6 and CRP levels. Together, these indices collectively point to innate immune activation and a myeloid-biased leukocyte profile and align with the broader concept of immunosenescence—an age-related remodeling of immunity characterized by chronic inflammatory load (“inflammaging”), hematopoietic myeloid skewing, reduced naïve and total lymphocyte resilience, accumulation of senescent or exhausted T-cell phenotypes, and impaired resolution pathways [67–69]. This Age-related immune dysregulation may promote nervous-system aging through inflammatory signaling at neuroimmune interfaces, including the BBB, meninges, and choroid plexus. Remodeling of these borders is proposed to alter the signals that reach the parenchyma and to impair CSF drainage and glymphatic waste clearance. In this framework, brain maintenance tracks immunological rather than chronological age [70]. In SCZ, this phenotype appears to manifest prematurely [68] and may be present even in first-episode and antipsychotic-naive patients [71–74]. Remarkably, in 22q11.2 deletion syndrome—the strongest known genetic risk factor for SCZ—accelerated T-cell maturation and senescence markers already appear in childhood, [67, 68, 75] and patients show similar profiles of blood inflammatory markers as our inflammatory subgroup of SCZ (increased levels of IL-6, CRP, and neutrophils to lymphocyte ratios) – particularly those individuals with 22q11.2 DS who also suffer from psychosis [76, 77]. Moreover, children with 22q11.2DS show a reduced DTI-ALPS index, and those who later develop psychotic symptoms do not exhibit the normative age-related increase [78]. Consistent with the concept of immunosenescence, proportions of neuroinflammatory phenotypes in post-mortem brain samples from SCZ patients assessed in this study resembled those from controls that were 20-30 years older. The overall percentage of 43% in our post-mortem cohort identified by an unbiased clustering method aggregated across microglia and astrocytes per donor closely matches findings from previous studies that utilized pre-selected panels of inflammatory markers and consistently recover a high-inflammation biotype comprising roughly 30–50% of patients as estimated by brain tissue [19, 20] or peripheral blood [79, 80]. Importantly, high inflammation cases are not merely a statistical tail of the same distribution: they show coordinated upregulation of innate cytokines and acute-phase transcripts [19, 20, 80], enrichment of macrophages/monocytes and adhesion-molecule signatures. Collectively, these findings suggest that systemic and neuroimmune aging are not independent phenomena but instead track the same latent axis. As further support for the relationship between immunological disturbance and glymphatic dysfunction in SCZ, we also observed a significant association between an immune-system-enriched polygenic risk score for SCZ and the DTI-ALPS index, connecting glymphatic impairment to the immunogenetic architecture of the disorder.

Like our post-mortem findings in the inflammatory subgroup of SCZ, AQP4 depolarization in acute inflammation is associated with increases in total AQP4 mRNA and protein through activation of the NF-κB pathway [81, 82]. Accordingly, several genes targeted by NF-κB and the interconnected JAK/STAT pathway were upregulated in parallel to AQP4 in our dataset (IL1B, CCL2, TLR2, JAK3, STAT3, see Figure 7 and Supplementary Table 2). Moreover, inflammation stimulates BMP2 signaling in astrocytes [83, 84] that was shown to both increase total AQP4 expression and dysregulate expression of key DAPC anchoring components (e.g. downregulation of DMD) leading to a transcriptional decoupling of AQP4 expression from its anchoring scaffold and – accordingly – a depolarized localization of AQP4 [85]. In a similar fashion, dysregulated expression of the AQP4-DAPC complex in our dataset was linked to astrocytic overexpression of different BMP pathway genes including BMPR1A and SMAD1 in the top 200 differentially expressed genes (Supplementary Table 1). Moreover, increased BMP signaling was shown to drive cytokine release likely through the NF-κB pathway in astrocytes in a mouse model of Parkinson’s disease [83]. In addition, BMP signaling has been identified as a key feature in experimental autoimmune encephalomyelitis [86] as well as iPSC-derived astrocytes from patients with SCZ [87]. Additional mechanisms of glymphatic dysfunction may include mechanical impediment of perivascular CSF flow by transmigrating leukocytes— supported by upregulation of endothelial genes involved in leukocyte diapedesis and downregulation of tight junction genes—as well as reduced apposition of astrocytic endfeet to the vasculature, as reported following systemic LPS administration [88]. A further mechanism may be the secretion of metalloproteases that cleave extracellular matrix proteoglycans, as suggested by the upregulation of ADAMTS9, ADAMTS9-AS1, and ADAMTS1 in endothelial cells, as well as ADAMTS9-AS2 and ADAMTS3 in astrocytes, in our dataset. While direct cleavage of β-dystroglycan (DAG1) by MMP-2/-9 has previously been demonstrated [89], secreted ADAMTS proteases could further destabilize the DAPC by remodeling its extracellular matrix environment.

Another pathway implicated by our in vivo and post-mortem data, that is closely linked to neuro-immunological and genetic models of SCZ, is mitochondrial dysfunction [90–92]. Mitochondrial integrity is essential for glymphatic function. As bioenergetic failure impairs both CSF production and perivascular clearance, an effect that may be further exacerbated by REM-sleep loss [93]. In idiopathic normal-pressure hydrocephalus (iNPH) and idiopathic intracranial hypertension (IIH), cortical biopsies reveal swollen mitochondria with disrupted cristae preferentially accumulated in perivascular astrocyte endfeet; the burden of these organelles correlates inversely with polarized AQP4 expression and directly with astrogliosis [94, 95]. ATP generated by oxidative phosphorylation drives choroid plexus Na⁺/K⁺-ATPase activity and astrocytic volume regulation, while mitochondrial ROS activate the NLRP3 inflammasome, amplifying microglial and secondary astrocytic dysfunction [96, 97]. Thus, energy failure and oxidative stress at both the choroid plexus and the astrocytic endfoot restrict CSF–ISF exchange. These mitochondrial abnormalities may feed a self-reinforcing inflammatory loop that intensifies with aging. An inflamed choroid plexus can lose checkpoint function: imbalanced T-cell cytokines, type-I interferon and complement, together with failure of local anti-inflammatory metabolic pathways, sustain choroid plexus and downstream brain inflammation with age [98]. Leakage of mtDNA activates cGAS– STING signalling, while ROS oxidize redox-sensitive transcription factors (NF-κB, AP-1, Nrf2), thereby sustaining cytokine production, mitochondrial damage [99–102], senescence and AQP4 depolarization [103–105]. Parallel bidirectional ROS– inflammation cycles have been documented in MDD and in SCZ-rat models, where mitochondrial gene alterations and redox imbalance generate circuit-level deficits [106, 107]. Because mitochondria are enriched within endfeet and support local Ca²⁺ buffering, their failure destabilizes the Ca²⁺-dependent signaling required for AQP4 polarization and vascular–glial coupling [108, 109]. The resulting loss of polarized water flux likely abolishes the osmotic driving forces for perivascular clearance.

Several constraints bound the interpretation of these findings. For our in vivo cohort, no blood samples were available from healthy control subjects. We therefore cannot determine whether imaging signatures of glymphatic dysfunction are specific to an inflammatory subgroup of SCZ or related to a transdiagnostic inflammatory immunosenescent phenotype that is enriched in SCZ. Findings from our post-mortem cohort rather support the latter hypothesis. Replication in other conditions with overlapping innate-immune signatures will be required before specificity can be claimed. Moreover, signatures of glymphatic dysfunction were derived from rather indirect measures (DTI-ALPS index, CPV, brain transcriptomics) and should be validated by brain proteomics and direct measurements of glymphatic flow. Intrathecal contrast would offer a more proximate readout but is difficult to justify at scale in this vulnerable patient population. However, the strong convergence of these MRI-derived proxies with independent post-mortem transcriptomic data indicates that the associations reported here are unlikely to represent artefacts of any single modality. Collectively, we propose that glymphatic dysfunction is embedded in a self-reinforcing cycle of inflammation, mitochondrial dysfunction, and oxidative stress, which together promote accelerated biological and brain aging. These interrelated processes are central to the pathophysiology of SCZ and likely extend to other neuropsychiatric disorders, highlighting their potential as transdiagnostic therapeutic targets to improve health outcomes beyond conventional diagnostic boundaries.

## 4. MATERIALS AND METHODS

### 4.1 Recruitment and clinical assessment

Structural MRI, diffusion tensor imaging (DTI) and neurocognitive data as well as blood samples for genotyping and inflammatory markers were collected within a clinical trial comparing different antipsychotic treatment strategies in SCZ (APIC, NCT02435095). For a total of 115 patients, both structural MRI and DTI data were available. The patient sample comprised both inpatients and outpatients, aged 18-65 years with a diagnosis of SCZ who were recruited from the Department of Psychiatry, Psychotherapy & Psychosomatics of RWTH Aachen University Hospital, as well as five affiliated academic hospitals (Alexianer Krankenhaus, Aachen; ViaNobis Gangelt; Klinik Königshof Krefeld; LVR Klinik Langenfeld and LVR Klinikum Düsseldorf). The primary inclusion criterion was a clinical diagnosis of SCZ, verified through the Structured Clinical Interview for DSM Disorders (SCID) based on DSM-5 (Diagnostic and Statistical Manual of Mental Disorders, Fifth Edition) criteria, conducted by trained psychiatrists. Participants also needed to be MRI-compatible and mentally as well as contractually capable of adhering to medical staff instructions. Exclusion criteria included significant somatic conditions that might influence study conduct (e.g., epilepsy, cancer and etc.), presence of cardiac pacemakers, body piercings or other metallic objects, pregnancy or lactation, prior inadequately documented antipsychotic treatment, court-or authority-mandated hospitalization, employment or dependency relationships with the sponsor or investigator, and concurrent involvement in another clinical study. Antipsychotic drugs taken by patients with SCZ were standardized to a haloperidol equivalent dose for each subject. Haloperidol equivalent doses were determined by dividing the daily dosage of each antipsychotic by its defined daily dose (DDD) and then multiplying this ratio by the DDD of haloperidol [110, 111]. Neurocognitive functioning was assessed using the Brief Assessment of Cognition in schizophrenia (BACS) [112]. For comparison purposes, we included structural MRI and DTI data from 73 healthy individuals participating in various study segments within the same research network (APIC), though laboratory or neurocognitive data were not available for this group. The study protocol received ethical approval from the North Rhine medical association’s ethics committee (AEKNO) and RWTH Aachen University Hospital’s local regulatory authority (EK 156/16). The study arms from which the healthy control subjects were derived also received approval from the RWTH Aachen University Hospital’s local regulatory authority (EK 226/15, EK 050/17, EK 188/17, EK 059/20).

Post-mortem sn-RNA-Seq data were obtained from two publicly available datasets [35, 113] comprising a total of 92 control donors and 95 donors with SCZ. Studies generating the post-mortem datasets received approval by their institutional review boards (22-0523, HE2018/351). Sociodemographic characteristics of the above data are presented in Supplementary Table 1.

### 4.2 Statistics

Statistical analyses were conducted in R software (version 4.2.3, https://www.r-project.org/) [114–116]. For most statistical analyses, linear models were employed that used age and gender as covariates. When outliers were present, robust linear models were used instead. Linear models were estimated with HC3 robust standard errors when unequal variance was identified. Overall group effects were assessed using F-tests for linear models and Wald-type tests for robust linear models or linear models with HC3 robust standard errors. Post hoc group comparisons were conducted using Dunnett’s tests for linear models and estimated marginal means contrasts with false discovery rate (FDR) correction of p-values for robust linear models or linear models with HC3 robust standard errors. Likelihood ratio tests (LRTs) were applied to compare nested linear regression models and to assess overall group effects on the regression relationship.

To compare group distributions of single variables, individual data points are overlaid on violin plots and combined with boxplots that indicate median and interquartile range (IQR). Points with error bars denote group means ± 95% CI for variables tested with two-sided hypotheses, whereas 90% CI intervals were used for variables tested with one-sided hypotheses. More details on each statistical test and post hoc comparisons are given in the respective figures as well as the corresponding paragraphs in the methods section.

### 4.3 MRI sequence parameters

MRI was performed using a 3 Tesla Tim Trio Scanner (Siemens, Erlangen, Germany) equipped with a 20-channel head coil. A T1-weighted 3D sequence (MPRAGE, echo time = 3.03 ms; repetition time = 2000 ms; inversion time = 900 ms; flip angle = 9°; field of view = 256×256 mm2; voxel size = 1×1×1 mm3; 176 sagittal partitions) was obtained for anatomical reference imaging. Diffusion Weighted Imaging data were collected using a spin-echo EPI sequence with the following parameters: repetition time = 7000 ms; echo time = 69 ms; field of view = 192×192 mm2; voxel size = 2×2×2 mm3; 65 slices; parallel imaging factor = 2; b value = 1000 s/mm2; 64 diffusion gradient directions.

### 4.4 CPV and DTI-ALPS index analysis

CPV was derived using two complementary approaches. First, CPV was calculated using a deep learning algorithm for automated segmentation of the T1-weighted structural MRI data [117] (https://github.com/hettk-/chp_seg). Second, CPV was independently quantified using a pipeline that combined atlas-based segmentation with a Gaussian mixture model (GMM): choroid plexus labels were initially obtained from FreeSurfer and subsequently refined using a GMM-based segmentation algorithm [24] (https://github.com/EhsanTadayon/choroid-plexus-segmentation). For both methods, individual CPV was normalized by the respective whole brain volume, used as an estimate of total intracranial volume (TIV), which was calculated using SIENA [118] of the FSL 6.0.1 toolbox (www.fmrib.ox.ac.uk/fsl). CPV measures from the two approaches were each z-standardized, and the mean of the two standardized values was used as the composite CPV index.

For DTI data, the default preprocessing pipeline according to FSL’s user guide was utilized (https://fsl.fmrib.ox.ac.uk/fsl/fslwiki/FDT/UserGuide), then Fractional Anisotropy (FA) and diffusivity maps in the X, Y and Z directions (*D_x_*, *D_y_*, *D_z_*) were generated. DTI-ALPS estimates glymphatic function by measuring CSF flow along the perivascular space. Due to the spatial orientation of vessels, at paraventricular projection and association fibers, the perivascular CSF flow is mainly reflected by diffusivity along the x-axis (*D_x_*) which is perpendicular to the diffusivity of the projection (*D_z_*) and association fibers (*D_y_*). *D_x_* is then normalized by the background diffusivity level which is reflected by *D_y_* for projections fibers and *D_z_* for association fibers.

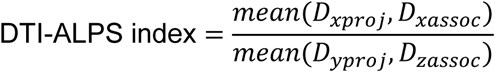

The corona radiata and superior longitudinal fasciculus were chosen as projection and association fibers, respectively, and automatically defined using FSL 6.0.1 toolbox. DTI-ALPS was quantified for each hemisphere as well as the bihemispheric average. Manual quality control was done as previously reported [23]. For a systematic comparison of DTI-ALPS index and CPV between patients with SCZ and healthy control subjects we employed linear mixed models considering group, gender, the total intracranial volume and age as independent between-subject variables and laterality as within-subject variable. To rule out medication effects, we performed subgroup analyses comparing DTI-ALPS index and CPV between patients with (SCZ AP+) or without antipsychotic treatment (SCZ AP-) and healthy controls. We applied linear models including group, age, total intracranial volume and gender as independent variables. To account for unequal variances, the linear models were estimated with HC3 robust standard errors. Overall group differences were assessed using Wald’s chi-square test. Estimated marginal means contrasts were used to compare the two SCZ subgroups with the CTR group, and false discovery rate (FDR) correction was applied to control for multiple comparisons.

### 4.5 Systemic inflammation and inflammatory markers

Whole blood counts of platelets, total leukocytes, monocytes, lymphocytes, and neutrophils, together with serum concentrations of IL-6 and CRP -- two common peripheral inflammatory markers – were obtained from 115 patients with SCZ. Blood samples from healthy controls were not available. The blood cell counts were used for the calculation of systemic inflammation markers, including PIV (platelets × neutrophils × monocytes / lymphocytes), SIRI (neutrophils × monocytes / lymphocytes), SII (platelets × neutrophils / lymphocytes), PLR (platelets / lymphocytes), MLR (monocytes / lymphocytes), and NLR (neutrophils / lymphocytes) according to previous papers [119, 120]. Blood cell counts were analyzed using a flow cytometer (Sysmex XN).

IL-6 concentrations were quantified through electrochemiluminescence immunoassay (ECLIA) using the Elecsys IL-6 kit on a Roche/Hitachi Cobas e411 analyzer. CRP levels were assessed utilizing the particle-enhanced immunoturbidimetric technique ACN 8217 on analyzer (Roche/Hitachi Cobas c701).

### 4.6 Genotyping and pathway related PRS analysis

To investigate the relationship between the glymphatic system and SCZ genetics, we studied pathway-related top hit polygenic risk scores (PRS) of SCZ based on DNA samples from our patients with SCZ. Thereto, an array-based genotyping was conducted using the Illumina Infinium Global Screening Array 24 v3.0 (Illumina, San Diego, CA, USA) and genotype information for each index SNP of the 287 loci with genome wide significance for SCZ according to the most recent GWAS [34] was retrieved.

Using gene ontology (GO) terms provided by the R biomaRt package [121], all genes which were mapped to the 287 loci by Trubetskoy [34] were assigned to different biological pathways related to the glymphatic system, namely “aging”, “immune system” or a “rest” category. To be assigned to the aging pathway, a gene’s GO terms had to include the strings “reactive oxygen”, “senescence” or “genome maintenance”. For the immune system pathway, respective GO terms had to include the strings “immune system” or “immune response”. If a gene did not match either pathway, it was assigned to the “rest category”. Some genes were assigned to both the “aging” and “immune system” pathways. Finally, pathway-related polygenic risk scores were calculated for each patient as the weighted sum of the genotype dosage of each pathway-related SNP applying the log-transformed odds ratio (OR) as the weighting factor:

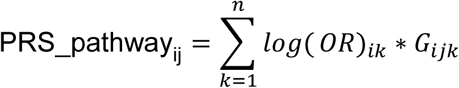

where PRS_pathwayij denotes the PRS of pathway i for person j, log(OR)ik denotes the log-transformed odds ratio of index SNP k for pathway i and Gijk is the allele dose of the minor allele of the respective SNP for person j.

### 4.7 Structural equation modeling (SEM)

To integrate inflammatory, glymphatic function, cognition dysfunction within a single explanatory framework, we specified a structural equation model (SEM) in the SCZ sample (n = 115). SEM was chosen over a series of separate regressions because it estimates all directional paths simultaneously and propagates the uncertainty of each path into the estimation of indirect effects, thereby allowing an explicit test of whether the association between peripheral inflammation and cognitive performance is transmitted through imaging markers of glymphatic function.

In the model, peripheral inflammation predicted glymphatic function, which in turn predicted cognitive performance. Peripheral inflammation was operationalized as a composite of the first and third principal components. PC1 was sign-inverted so that higher values of both components indicate greater inflammatory load; the two components were then z-standardized, averaged, and the average was z-standardized again to form the composite inflammatory index. The bihemispheric DTI-ALPS index and the composite CPV were modelled as mediators, and the BACS total score as the distal outcome.

All model variables were z-standardized prior to model fitting, so that path coefficients can be interpreted as fully standardized effects. The model was estimated using the R lavaan package (version 0.6-21). Global model fit was assessed using the model chi-square statistic, the comparative fit index (CFI), the Tucker-Lewis index (TLI), the root mean square error of approximation (RMSEA), and the standardized root mean square residual (SRMR). Fit was considered acceptable for CFI and TLI ≥ 0.95, RMSEA ≤ 0.06 and SRMR ≤ 0.08. Competing nested models were compared using the likelihood-ratio chi-square difference test.

Because the direction of each path had been hypothesized a priori — a higher inflammatory load being associated with poorer glymphatic function, and poorer glymphatic function in turn with poorer cognitive performance — the p-values reported for the respective path coefficients are one-tailed.

### 4.8 Single nucleus – RNA-Seq data analysis

#### 4.8.1 Download and quality control of snRNA-Seq public datasets

Preprocessed single-nuclei count data from Fröhlich et al. [54] were obtained from GEO -Gene Expression Omnibus (accession number GSE254569). Preprocessed, demultiplexed single nuclei count data for the HBCC and MSSM cohorts, part of the PsychAD consortium [35], were obtained from the synapse repository syn68559919. Cell type annotations for the HBCC and MSSM cohorts were used as provided in the publication. For Fröhlich et al., cell types were iteratively cleaned and re-annotated in order to remove doublets and ensure consistent cell type annotations across the cohorts. Single nuclei annotated to different general cell types (microglia, astrocytes, endothelial cells, excitatory neurons, inhibitory neurons, and oligodendrocytes with OPCs) were subset into cell-type specific objects and further analyzed using scanpy [122]. Within each cell-type specific objects, normalization was performed, the top variable genes were detected, the normalized counts were scaled, and PCA was performed followed by harmony correction [123] treating lib batch and Sex as covariates. Leiden clustering [124] was performed on each of the objects and clusters co-expressing canonical marker genes of multiple different cell types were filtered. The remaining cells were re-integrated, clustered, and annotated according to the relative expression of cell-type specific marker genes. Using the refined cell type annotations from Fröhlich et al. and the annotations provided for the HBCC and MSSM cohorts, pseudobulks were generated for each cell type by aggregating cells within each donor annotated to the same cell type and summing the counts for each gene across all cells with a minimum cutoff of 10 cells per pseudobulk sample.

#### 4.8.2. Filtering samples by diagnosis and comorbidities

Due to the high degree of heterogeneity of samples in the MSSM and HBCC cohorts with respect to various comorbidities and clinical diagnoses, stringent filtering of samples was applied to remove potential covariates that would impact downstream interpretation of results. Donors with any neurodegenerative disease including Alzheimer’s disease, Parkinson’s disease, multiple sclerosis, dementia with Lewy bodies, cerebral atrophy, tauopathy, frontotemporal dementia, dementia, MCI (mild cognitive impairment), as well as donors with Alzheimer’s Braak stage at least 4 or CERAD score at least 3, neurological disorders including epilepsy or seizures, or other neuropsychiatric disorders including major depressive disorder and bipolar disorder, were removed from downstream analyses. The remaining donors were then classified as SCZ or controls according to the metadata provided in the publication. For the Fröhlich et al. cohort, donors with a diagnosis of bipolar disorder, schizoaffective disorder or major depression were filtered from downstream analyses.

#### 4.8.3. Joint clustering of microglia and astrocytes into high and low inflammation groups

Microglia and astrocyte pseudobulk objects were integrated together by concatenating the expression profiles from each donor containing both a microglia and astrocyte pseudobulk object (donors with at least 10 cells of each cell type). Due to the high discrepancy in age between controls and SCZ donors in the HBCC and MSSM cohorts, age-matching was subsequently performed within these two cohorts separately by randomly sampling controls, enforcing a similar age distribution across controls and SCZ donors. The microglia and astrocyte pseudobulk objects were analyzed using Seurat [125]. Microglia and astrocyte pseudobulk objects were normalized separately with a normalization factor of 10,000 and the top 2,000 variable genes were detected. These objects were then concatenated into a single Seurat object with each sample comprised of the joint expression profiles of highly variable microglia and astrocyte genes. The normalized counts were scaled, and the samples were integrated with the IntegrateLayers() function in Seurat using canonical correlation analysis (CCA), treating the cohorts (MSSM, HBCC and Fröhlich) as separate batches in the data. A k-nearest neighbor graph was then calculated using the first 15 dimensions and 20 nearest neighbors from the integrated CCA dimension reduction as well as a UMAP dimension reduction. Clusters were then measured using the Louvain clustering algorithm [126] with a resolution of 0.2 with the FindClusters() function in Seurat. This resulted in two clusters which were annotated as high and low inflammation based on the relative expression of homeostatic and inflammatory marker genes in microglia and astrocytes.

#### 4.8.4 Logistic regression inflammation group subtypes

Logistic regression analysis was performed to model the probability of a sample belonging to the high (1) or low (0) inflammation group based on different independent variables using the glm() function in R. Two models were tested, including Disease (SCZ or CTR), Age and Sex as independent variables: glm(immune_subtype ∼ Disease + Age + Sex, data = metadata, family = “binomial”), and in addition including pH, PMI (post-mortem interval), and Source (Fröhlich, HBCC or MSSM) as independent variable: glm(immune_subtype ∼ Disease + Age + Sex + pH + PMI + Source, data = metadata, family = “binomial”). Likelihood ratio tests were subsequently performed to determine if a model with a combination of interaction terms between Disease, Age and Sex significantly improved the model using an Anova test with the anova() function in R.

#### 4.8.5. Glymphatic dysfunction signature

A glymphatic dysfunction signature score was calculated for each astrocyte pseudobulk sample to summarize changes observed in genes involved in the glymphatic system in SCZ patients within the high inflammation subgroup. Specifically, we summarized the previously observed significant upregulation of AQP4 and DTNA and downregulation of SNTA1 and DMD in SCZ into a common score. First, the expression matrix was corrected for dataset source using the combat function from the sva package (v.3.46.0) [127] in R. Subsequently, R package singscore v. 1.18.0 [128] was used to rank the batch-corrected expression matrix (function rankGenes) and to calculate signature scores using the function simpleScore(upSet = c(’AQP4’,’DTNA’), downSet = c(’SNTA1’,’DMD’)).

#### 4.8.6. Aging signature

Aging signatures for microglia and astrocytes were obtained from Fröhlich et al. who derived cell type-specific aging signatures by identifying genes whose expression was associated with donor age within each annotated cell type after controlling for technical and biological covariates [54]. The signature list was filtered for assay ‘Astrocytes’ or ‘Microglia’ and genes with adj.P.Val < 0.05 were selected. Signature scores were calculated for each astrocyte or microglia pseudobulk sample using the singscore v. 1.18.0 [128]. The batch-corrected expression matrix was used to calculate a rank matrix using rankGenes(). Subsequently, the simpleScore() function was used to calculate a signature score specifying genes with logFC > 0 in the aging signature as ‘upSet’ and those with logFC < 0 as ‘downSet’.

#### 4.8.7. DEG Analysis

Genes were tested for differential expression using uncorrected normalized expression values and linear models: lm(Expression ∼ Diagnosis + Age + Sex + PMI + pH + Dataset_source). Subsequently, a type II F-test was applied using Anova(type = 2) from the R car package [129] with options(contrasts = “contr.sum”) and p-values were corrected for multiple testing using p.adjust(method = ‘fdr’). Linear models with and without interaction terms between Diagnosis, age and sex were compared using a likelihood-ratio test implemented in the anova() function. Normalized pseudobulk expression profiles of astrocytes, microglia and endothelial cells were separately corrected for batch effects driven by different cohorts (MSSM, HBCC and Fröhlich) using the ComBat method [130] with the function ComBat() in the R package sva (v.3.46.0) [127]. Pearson correlation coefficients were estimated between the glymphatic dysfunction score and gene expression profiles in each cell type, excluding genes with less than 10 counts across all pseudobulk samples. Corrected expression values were subsequently scaled using the ScaleData() function in Seurat.

### 4.9 Brain-Age Modeling

Brain age gap estimates were derived using support vector regression (SVR) models trained to predict chronological age from whole-brain regional grey matter and CSF volumes obtained via voxel-based morphometry (VBM) of T1 images. VBM was performed using v7771 software (FIL Methods Group, University College London, London, UK) implemented in MATLAB R2020b v9.9 (The MathWorks, Natick, MA, USA). VBM preprocessing was performed using the CAT12(12.7)-toolbox in default mode [131]. Following brain extraction, a registration process was applied to ensure correct voxel positions with regard to the anatomical locations. A spatial transformation was performed through affine registration aligning the image to the Montreal Neurological Institute (MNI) space. Affine registration features three rotations, three translations, three scalings and three skew/shears along 12 degrees of freedom in 3-dimensional space. Linear registration is necessary to remove movement artefacts. It also initializes non-linear registration, which is indispensable for between-subject group comparisons. In the transformation, a 4×4 matrix is multiplied by the coordinate vectors for defining the “warped” coordinates in the deformation field. The CAT toolbox uses the geodesic approach for registration, which searches for the shortest path between the coordinates of two brain images in a complex high-dimensional space [132]. This method expands on affine registration by allowing more nuanced, non-linear transformations. Differences between the images are expressed through a cost function, which the algorithm aims to minimize. Bias due to radio frequency (RF) coil uniformity, eddy currents or the anatomy of the measured subject may occur [133]. After registration, the CAT toolbox uses the unified segmentation approach, in which higher computing costs are exchanged for better accuracy [134]. In this approach bias correction, registration through non-linear warping, and segmentation are done in a single step. This is supported by tissue probability maps in which an a priori likelihood for the presence of a particular tissue at a given voxel is delineated from existing neuroimaging data. Parcellation was done according to the Neuromorphometrics Atlas in which more than 100 high-resolution MRIs were manually labelled with high precision using established protocols and registered in the MNI-space [135]. The segmented images were visually inspected for non-brain structures by overlaying GM segments on the structural image for each subject. Additionally, the CAT toolbox provided the automated image and preprocessing quality scores for each subject. Image quality before preprocessing was assessed via a weighted overall image quality score accounting for resolution, noise and bias.

Standardization and model training were performed as follows: For each analysis, regional gray matter volumes and CSF volumes were scaled using a standard (z-score) scaler as implemented in scikit-learn, with the scaler choice held fixed within a given model. An ε-insensitive support vector regression (SVR) model with radial basis function (RBF) kernel was then fitted using the scikit-learn SVR implementation. Hyperparameters were optimized via grid search over the regularization parameter C, kernel width γ, and epsilon insensitive margin evaluated with five-fold cross validation, and the configuration achieving the lowest cross-validated mean absolute error was selected as the final estimator.

To enhance generalizability, the age-prediction model for healthy controls was trained on the combined sample of local healthy controls and a large, publicly available lifespan dataset (openBHB) [55]. The local control cohort was partitioned into five subsets; in each fold, one subset served as an out-of-fold test set while the remaining local controls, together with all openBHB controls, were used for training. Chronological age was then regressed onto the resulting out-of-fold predicted ages within the full control sample using simple linear regression, yielding a global age-bias correction model in line with established procedures [136]. For each control, the corrected predicted age was defined as the residual of this regression added back to the observed chronological age, and the corresponding brain age gap was obtained as the residual itself. The final SVR model with the tuned hyperparameters was subsequently refit to all control data and applied once to each patient to obtain raw predicted age, and the control-derived bias-correction regression was then used to compute each patient’s brain age gap as the residual of raw predicted age after adjusting for chronological age.

To evaluate the effect of glymphatic dysfunction on brain age gap, patients with SCZ were stratified into subgroups based on a median split of either the DTI-ALPS index or CPV. Individual brain age gap values were then analyzed using a linear model including age, sex, and group membership ([CTR, ALPS+, ALPS−] or [CTR, CPV−, CPV+]) as predictors. Overall group differences were assessed using F-tests, followed by Dunnett’s tests for pairwise comparisons against the control group. Given the a priori hypothesis of an increased brain age gap in patients relative to controls, one-sided (right-tailed) p-values were applied.

### 4.10 DNA methylation age

DNAm profiling of patients’ whole blood samples was performed using Illumina Infinium Methylation EPIC BeadChip arrays following the manufacturer’s recommended protocol (Illumina Inc., San Diego, CA, USA). Raw IDAT files from Illumina EPIC BeadChip arrays were processed in R using established pipelines for background correction, dye-bias adjustment, and probe filtering [137, 138]. Samples with poor probe detection and probes failing detection or known to overlap common SNPs or cross-reactive sites were excluded.

Epigenetic clocks were then computed for the patient cohort using the ENmix methscore framework with reference-based normalisation and imputation of missing CpG values [137]. From these, we derived the principal component–based GrimAge (PCGrimAge) as our primary DNAm age measure. GrimAge is a machine-learning– derived epigenetic clock that combines DNA-methylation surrogates of mortality-related plasma proteins and smoking exposure in a time-to-event model, providing an age-like estimate strongly associated with lifespan and healthspan [56, 139].

As an external reference, we used publicly available Illumina EPIC blood methylation data from GEO series GSE147740 (1024 healthy controls) [57], retaining only samples with valid age and sex information. Starting from the study’s quantile-normalised beta matrix, we reduced the data to CpGs required for PCGrimAge and applied k-nearest-neighbour imputation for residual missing values to obtain a control beta matrix aligned with the corresponding phenotype data.

Because the full methscore pipeline was not feasible on this large external dataset, we implemented a streamlined PCGrimAge calculation that reproduces ENmix’s internal PC-clock logic: applying the published GrimAge PC-clock model to the reduced control beta matrix to obtain PCGrimAge values for all controls.

We defined age-calibrated epigenetic age gaps by comparing observed PC-based clocks to the values expected in healthy controls of the same age. Using the external control dataset, we performed 5-fold cross-fitting for PCGrimAge. The controls were randomly partitioned into five folds. For each fold, we fit linear regression models in the training subset modelling PCGrimAge as a function of age.

The resulting models were then used to generate out-of-fold predictions for PCGrimAge in the held-out control subset of each fold. For the patients, the whole control dataset was used for training. The age gap for a given clock was defined as the difference between observed and expected PC-based age:

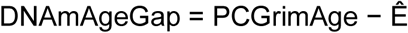

Positive age gaps indicate epigenetic ages higher than expected given chronological age and the control-based calibration, whereas negative gaps indicate lower-than-expected epigenetic ages. To assess the impact of glymphatic dysfunction on DNAm age gap, subgroups of SCZ were defined based on a median split of DTI-ALPS index or CPV and each subject’s age gap entered a linear model using age, sex and group ([CTR, ALPS+, ALPS-] or [CTR, CPV-, CPV+]). Group differences were assessed using F-tests and Dunnett’s tests were used for pairwise comparisons. Since we expected an increase of DNAm age gap in the patients, right-sided p-values were applied for Dunnett’s test.

## Data availability

Data supporting the findings of this study are available from the corresponding author upon reasonable request.

## Acknowledgements

The authors thank Laura Bell of the Audiovisual Media Center, Faculty of Medicine, RWTH Aachen University, for her excellent assistance with the design of the figures.

The authors thank Manuela Das Gupta for her contribution to blood sampling, and Dr. Michelle Schlingensief, as well as Jasmin Mühlenberg for their organizational help. Furthermore, they thank Cordula Kemper and the brain imaging facility of the interdisciplinary center for clinical research (IZKF Aachen) within the Medical Faculty of the RWTH Aachen University for their contribution to recording and storing MRI data. Moreover, the authors thank the other members of the APIC Consortium for their contribution to the recruitment of participants and organizational help: Marc Augustin,; Joachim Cordes; Emir Demirel; Thomas Dielentheis; Jan Dreher; Patrick Eisner; Michelle Finner-Prével; Gerhard Gründer; Frederik Hendricks; Jana Hovancakova; Peter Kaleta; Fatih Keskin; Miriam Kirchner; André Kirner-Veselinovic; Sarah Lammertz; Christina Lange; Federico Maria Larcher; Laura M. Lenzen; Eva Meisenzahl-Lechner; Jutta Muysers; Andrea Neff; Michael Plum; Erik Röcher; Axel Ruttmann; Sabrina Schaffrath; Frank Schneider; Georgios Schoretsanitis; Lara Schwemmer; Eva Stormanns; Antje Trauzeddel; Lina Winkler.

Use of artificial intelligence: Perplexity Pro was used to assist with literature searches, language editing, and the drafting and debugging of code. All AI-assisted outputs, including literature suggestions and code, were independently reviewed, verified, and, where appropriate, revised by the authors. The authors take full responsibility for the manuscript and all analyses.

## Author contributions

XT: study design, data analysis, interpretation of results, manuscript writing

MS: data analysis, interpretation of results, manuscript correction

DR: data analysis, interpretation of results, manuscript correction

MFi: data analysis, manuscript correction

DRG: interpretation of results, manuscript correction

MFu: data analysis, interpretation of results, manuscript correction

RA: data analysis, manuscript correction

SHH: data analysis, manuscript correction

KB: interpretation of results, manuscript correction GS: data analysis, manuscript correction

TF: resource allocation, interpretation of results, manuscript correction

KM: resource allocation, interpretation of results, manuscript correction

AJG: study design, data collection, data analysis, interpretation of results, manuscript writing

## Supplementary Figures

**Supplementary Figure 1.**
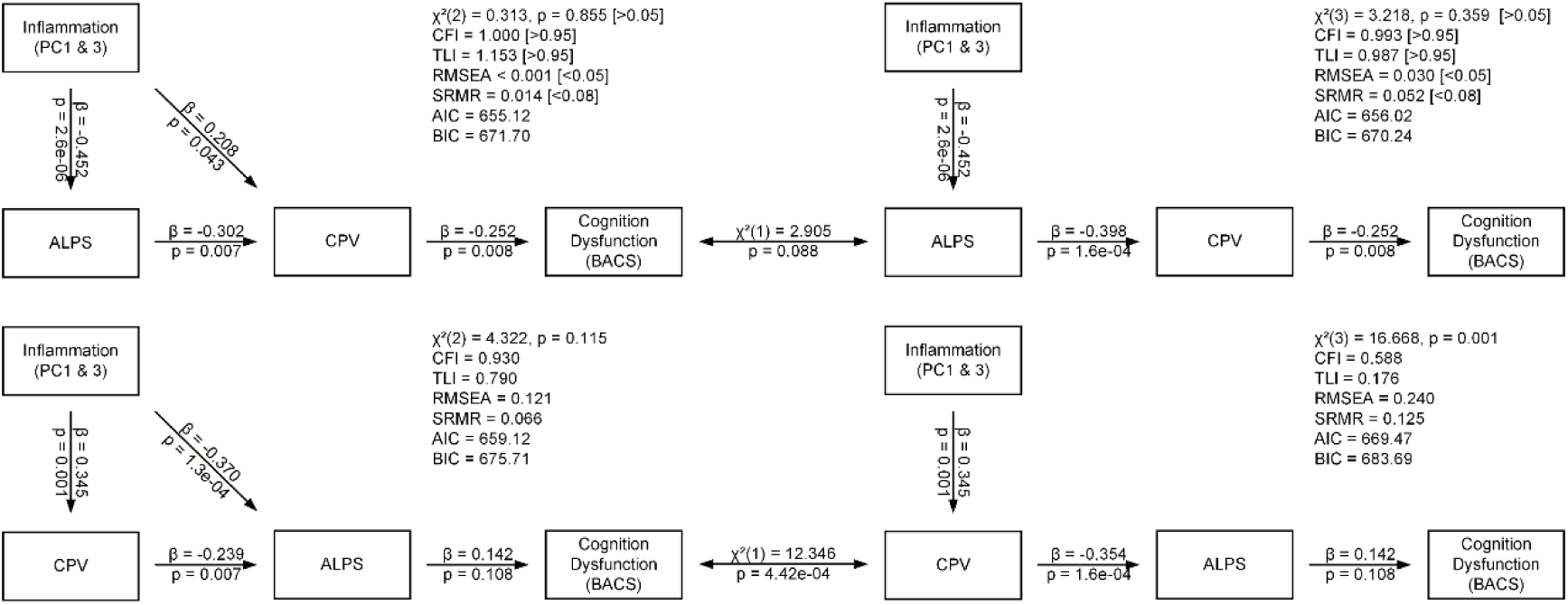
Comparison of four competing structural equation models. The two rows contrast the order of the two intermediate variables: in the upper row inflammation is related to cognition via the DTI-ALPS index and subsequently CPV, whereas in the lower row this order is reversed. The two columns contrast the specification of the direct path from inflammation to the second intermediate variable, which is freely estimated in the left column and fixed to zero in the right column. Double-headed arrows between the left and right panels of each row denote likelihood-ratio tests of this nested constraint. Conventional thresholds for acceptable model fit are given in brackets.

**Supplementary Figure 2.**
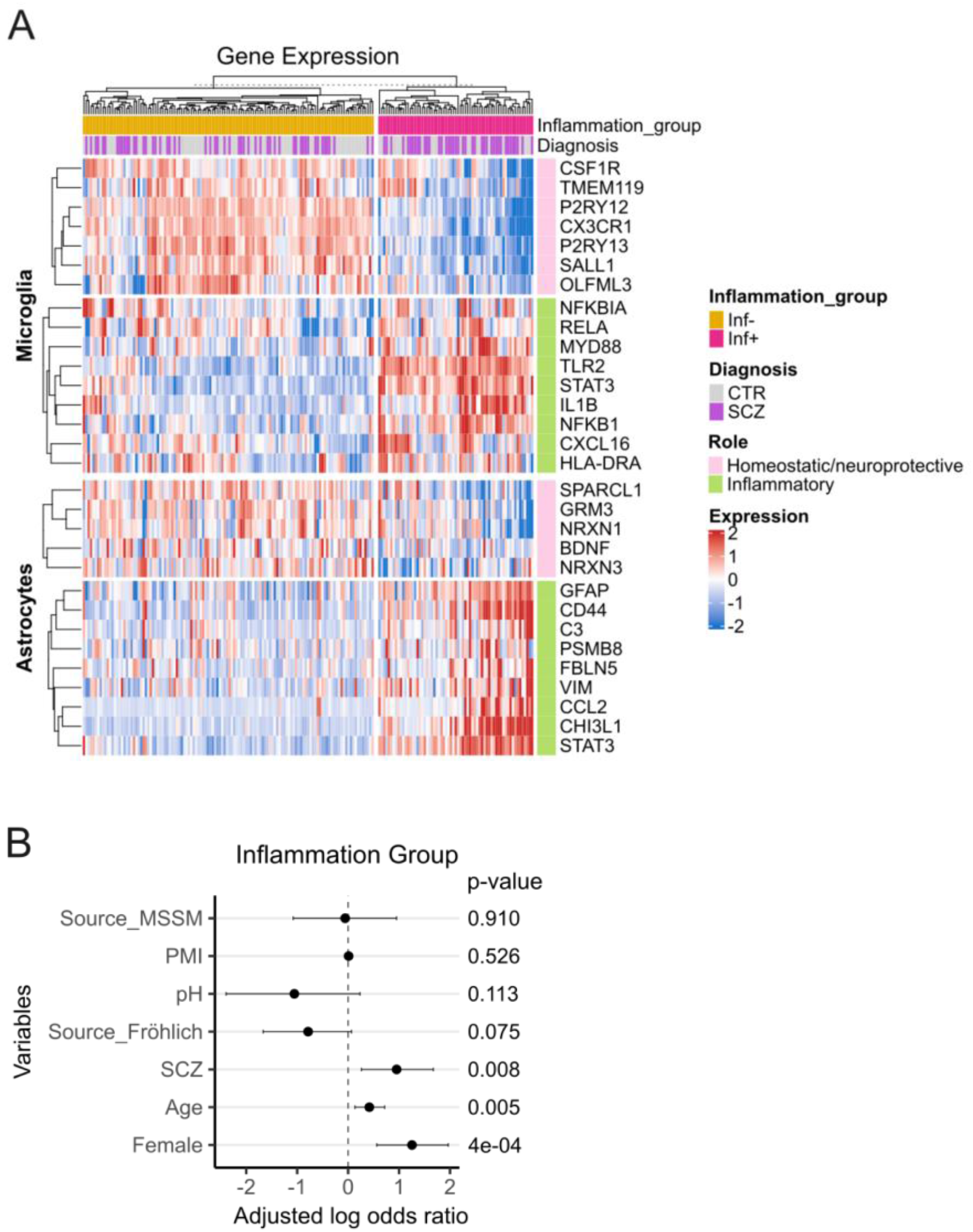
Inflammatory glial transcriptional signatures in separate donors into inflammation subgroups. **(A)** Heatmap showing normalized expression (z-score) of selected microglial and astrocytic genes across all samples. Columns represent individual samples and rows represent genes. Samples are annotated by inflammation group (Inf−, yellow; Inf+, magenta) and diagnosis (control (CTR), grey; schizophrenia (SCZ), purple) and hierarchically clustered within each inflammation group. Gene expression of homeostatic/neuroprotective (pink row annotation) and inflammatory (green row annotation) markers is shown for microglia and astrocytes. (B) Multivariable logistic regression analysis of factors associated with assignment to the Inf+ subgroup. Dataset source, diagnosis, gender, age, pH and PMI were included as explanatory variables in the model. Points indicate adjusted log-odds ratios, and lines represent 95% confidence intervals. Coefficient P values are indicated.

## References

1. Hjorthøj, C., et al., Years of potential life lost and life expectancy in schizophrenia: a systematic review and meta-analysis. Lancet Psychiatry, 2017. 4(4): p. 295–301.

2. McCutcheon, R.A., R.S.E. Keefe, and P.K. McGuire, Cognitive impairment in schizophrenia: aetiology, pathophysiology, and treatment. Mol Psychiatry, 2023. 28(5): p. 1902–1918.

3. Gupta, S. and P. Kulhara, What is schizophrenia: A neurodevelopmental or neurodegenerative disorder or a combination of both? A critical analysis. Indian J Psychiatry, 2010. 52(1): p. 21–7.

4. Falkai, P. and A. Schmitt, Failed regeneration and inflammation in schizophrenia: two sides of the same coin? J Neural Transm (Vienna), 2022. 129(5-6): p. 611–615.

5. Constantinides, C., et al., Brain ageing in schizophrenia: evidence from 26 international cohorts via the ENIGMA Schizophrenia consortium. Mol Psychiatry, 2023. 28(3): p. 1201–1209.

6. Koutsouleris, N., et al., Exploring Links Between Psychosis and Frontotemporal Dementia Using Multimodal Machine Learning: Dementia Praecox Revisited. JAMA Psychiatry, 2022. 79(9): p. 907–919.

7. Yu, W.Y., et al., Short telomeres in patients with chronic schizophrenia who show a poor response to treatment. J Psychiatry Neurosci, 2008. 33(3): p. 244–7.

8. Gurvich, C., et al., The relationship between cognitive clusters and telomere length in bipolar-schizophrenia spectrum disorders. Psychol Med, 2023. 53(11): p. 5119–5126.

9. Shirai, T., et al., Meta-analyses of epigenetic age acceleration and GrimAge components of schizophrenia or first-episode psychosis. Schizophrenia (Heidelb), 2024. 10(1): p. 108.

10. Ori, A.P.S., et al., Meta-analysis of epigenetic aging in schizophrenia reveals multifaceted relationships with age, sex, illness duration, and polygenic risk. Clin Epigenetics, 2024. 16(1): p. 53.

11. Fizjková, I., J. Dragašek, and P. Račǎy, Mitochondrial Dysfunction, Altered Mitochondrial Oxygen, and Energy Metabolism Associated with the Pathogenesis of Schizophrenia. Int J Mol Sci, 2023. 24(9).

12. Okusaga, O.O., Accelerated aging in schizophrenia patients: the potential role of oxidative stress. Aging Dis, 2014. 5(4): p. 256–62.

13. Dudeck, L., et al., Differences in Blood Leukocyte Subpopulations in Schizophrenia: A Systematic Review and Meta-Analysis. JAMA Psychiatry, 2025. 82(5): p. 492–504.

14. Fillman, S.G., et al., Elevated peripheral cytokines characterize a subgroup of people with schizophrenia displaying poor verbal fluency and reduced Broca’s area volume. Mol Psychiatry, 2016. 21(8): p. 1090–8.

15. Trépanier, M.O., et al., Postmortem evidence of cerebral inflammation in schizophrenia: a systematic review. Mol Psychiatry, 2016. 21(8): p. 1009–26.

16. North, H.F., et al., A schizophrenia subgroup with elevated inflammation displays reduced microglia, increased peripheral immune cell and altered neurogenesis marker gene expression in the subependymal zone. Transl Psychiatry, 2021. 11(1): p. 635.

17. Gober, R., et al., Increased inflammasome protein expression identified in microglia from postmortem brains with schizophrenia. J Neuropathol Exp Neurol, 2024. 83(11): p. 951–966.

18. Gober, R., et al., Microglia activation in postmortem brains with schizophrenia demonstrates distinct morphological changes between brain regions. Brain Pathol, 2022. 32(1): p. e13003.

19. Fillman, S.G., et al., Markers of inflammation and stress distinguish subsets of individuals with schizophrenia and bipolar disorder. Transl Psychiatry, 2014. 4(2): p. e365.

20. Fillman, S.G., et al., Increased inflammatory markers identified in the dorsolateral prefrontal cortex of individuals with schizophrenia. Mol Psychiatry, 2013. 18(2): p. 206–14.

21. Iliff, J.J., et al., A paravascular pathway facilitates CSF flow through the brain parenchyma and the clearance of interstitial solutes, including amyloid β. Sci Transl Med, 2012. 4(147): p. 147ra111.

22. Mogensen, F.L., C. Delle, and M. Nedergaard, The Glymphatic System (En)during Inflammation. Int J Mol Sci, 2021. 22(14).

23. Taoka, T., et al., Evaluation of glymphatic system activity with the diffusion MR technique: diffusion tensor image analysis along the perivascular space (DTI-ALPS) in Alzheimer’s disease cases. Jpn J Radiol, 2017. 35(4): p. 172–178.

24. Tadayon, E., et al., Improving Choroid Plexus Segmentation in the Healthy and Diseased Brain: Relevance for Tau-PET Imaging in Dementia. J Alzheimers Dis, 2020. 74(4): p. 1057–1068.

25. Zhou, Y.F., et al., Choroid Plexus Enlargement and Allostatic Load in Schizophrenia. Schizophr Bull, 2020. 46(3): p. 722–731.

26. Lizano, P., et al., Association of Choroid Plexus Enlargement With Cognitive, Inflammatory, and Structural Phenotypes Across the Psychosis Spectrum. Am J Psychiatry, 2019. 176(7): p. 564–572.

27. Yu, L., et al., The relationship between choroid plexus volume, DTI-ALPS index, kidney function, and amyloid pathology in alzheimer’s disease. Neuroradiology, 2025. 67(12): p. 3587–3597.

28. Takahashi, T., et al., Longitudinal changes in DTI-ALPS and choroid plexus volume relative to CSF biomarkers during lecanemab treatment in mild cognitive impairment: A pilot study. Alzheimers Dement (Amst), 2026. 18(3): p. e70375.

29. Zhao, R., et al., Potentiating Cerebral Perfusion Normalizes Glymphatic Dynamics in Systemic Inflammation. Adv Sci (Weinh), 2025. 12(47): p. e03576.

30. Soden, P.A., A.R. Henderson, and E. Lee, A Microfluidic Model of AQP4 Polarization Dynamics and Fluid Transport in the Healthy and Inflamed Human Brain: The First Step Towards Glymphatics-on-a-Chip. Adv Biol (Weinh), 2022. 6(12): p. e2200027.

31. Lin, R., et al., Association of glymphatic system function with peripheral inflammation and motor symptoms in Parkinson’s disease. NPJ Parkinsons Dis, 2025. 11(1): p. 62.

32. Liang, Q., et al., Inflammation and psychomotor retardation in depression: The moderating role of glymphatic system. Brain Behav Immun, 2025. 127: p. 387–395.

33. Borovcanin, M.M., et al., Interleukin-6 in Schizophrenia-Is There a Therapeutic Relevance? Front Psychiatry, 2017. 8: p. 221.

34. Trubetskoy, V., et al., Mapping genomic loci implicates genes and synaptic biology in schizophrenia. Nature, 2022. 604(7906): p. 502–508.

35. Fullard, J.F., et al., Population-scale cross-disorder atlas of the human prefrontal cortex at single-cell resolution. Sci Data, 2025. 12(1): p. 954.

36. Campeau, A., et al., Multi-omics of human plasma reveals molecular features of dysregulated inflammation and accelerated aging in schizophrenia. Mol Psychiatry, 2022. 27(2): p. 1217–1225.

37. Bojarskaite, L., et al., Role of aquaporin-4 polarization in extracellular solute clearance. Fluids Barriers CNS, 2024. 21(1): p. 28.

38. Neely, J.D., et al., Syntrophin-dependent expression and localization of Aquaporin-4 water channel protein. Proc Natl Acad Sci U S A, 2001. 98(24): p. 14108–13.

39. Martins-Ferreira, R., et al., The Human Microglia Atlas (HuMicA) unravels changes in disease-associated microglia subsets across neurodegenerative conditions. Nat Commun, 2025. 16(1): p. 739.

40. Liu, H., et al., Fibroblast Growth Factor 8 Suppresses Neurotoxic Astrocytes and Alleviates Neuropathic Pain via Spinal FGFR3 Signaling. Neurosci Bull, 2025. 41(12): p. 2218–2232.

41. Saba, J., et al., Astrocyte truncated tropomyosin receptor kinase B mediates brain-derived neurotrophic factor anti-apoptotic effect leading to neuroprotection. J Neurochem, 2018. 146(6): p. 686–702.

42. Butovsky, O. and H.L. Weiner, Microglial signatures and their role in health and disease. Nat Rev Neurosci, 2018. 19(10): p. 622–635.

43. Zhu, C., et al., Expression site of P2RY12 in residential microglial cells in astrocytomas correlates with M1 and M2 marker expression and tumor grade. Acta Neuropathol Commun, 2017. 5(1): p. 4.

44. Yuan, J.P., et al., Paeoniflorin regulates RhoA/ROCK1 and Nrf2 pathways in PDLIM1-dependent or independent manners in oxidative stressed melanocytes. Arch Dermatol Res, 2024. 316(7): p. 401.

45. Keeney, M.T., et al., LRRK2 regulates production of reactive oxygen species in cell and animal models of Parkinson’s disease. Sci Transl Med, 2024. 16(767): p. eadl3438.

46. Saito, Y., Selenoprotein P as an in vivo redox regulator: disorders related to its deficiency and excess. J Clin Biochem Nutr, 2020. 66(1): p. 1–7.

47. Hanmer, K.L. and D. Mavri-Damelin, Peroxidasin is a novel target of the redox-sensitive transcription factor Nrf2. Gene, 2018. 674: p. 104–114.

48. Barodia, S.K., R.B. Creed, and M.S. Goldberg, Parkin and PINK1 functions in oxidative stress and neurodegeneration. Brain Res Bull, 2017. 133: p. 51–59.

49. Murphy, C.E., A.K. Walker, and C.S. Weickert, Neuroinflammation in schizophrenia: the role of nuclear factor kappa B. Transl Psychiatry, 2021. 11(1): p. 528.

50. Yu, H., D. Pardoll, and R. Jove, STATs in cancer inflammation and immunity: a leading role for STAT3. Nat Rev Cancer, 2009. 9(11): p. 798–809.

51. Chang, J., et al., Gpr124 is essential for blood-brain barrier integrity in central nervous system disease. Nat Med, 2017. 23(4): p. 450–460.

52. Sohet, F., et al., LSR/angulin-1 is a tricellular tight junction protein involved in blood-brain barrier formation. J Cell Biol, 2015. 208(6): p. 703–11.

53. Wimmer, I., et al., PECAM-1 Stabilizes Blood-Brain Barrier Integrity and Favors Paracellular T-Cell Diapedesis Across the Blood-Brain Barrier During Neuroinflammation. Front Immunol, 2019. 10: p. 711.

54. Fröhlich, A.S., et al., Single-nucleus transcriptomic profiling of human orbitofrontal cortex reveals convergent effects of aging and psychiatric disease. Nat Neurosci, 2024. 27(10): p. 2021–2032.

55. Dufumier, B., et al., OpenBHB: a Large-Scale Multi-Site Brain MRI Data-set for Age Prediction and Debiasing. Neuroimage, 2022. 263: p. 119637.

56. Lu, A.T., et al., DNA methylation GrimAge strongly predicts lifespan and healthspan. Aging (Albany NY), 2019. 11(2): p. 303–327.

57. Robinson, O., et al., Determinants of accelerated metabolomic and epigenetic aging in a UK cohort. Aging Cell, 2020. 19(6): p. e13149.

58. Chen, T., et al., The brain washing system in sepsis-associated encephalopathy. J Neuroinflammation, 2025. 22(1): p. 277.

59. Erickson, M.A., et al., Lipopolysaccharide impairs amyloid β efflux from brain: altered vascular sequestration, cerebrospinal fluid reabsorption, peripheral clearance and transporter function at the blood-brain barrier. J Neuroinflammation, 2012. 9: p. 150.

60. Howes, O.D. and E.C. Onwordi, The synaptic hypothesis of schizophrenia version III: a master mechanism. Mol Psychiatry, 2023. 28(5): p. 1843–1856.

61. Khandaker, G.M., et al., Childhood infection and adult schizophrenia: a meta-analysis of population-based studies. Schizophr Res, 2012. 139(1-3): p. 161–8.

62. Sekar, A., et al., Schizophrenia risk from complex variation of complement component 4. Nature, 2016. 530(7589): p. 177–83.

63. Halstead, S., et al., Alteration patterns of peripheral concentrations of cytokines and associated inflammatory proteins in acute and chronic stages of schizophrenia: a systematic review and network meta-analysis. Lancet Psychiatry, 2023. 10(4): p. 260–271.

64. Zhu, Y., et al., Distinct Phenotypes of Inflammation Associated Macrophages and Microglia in the Prefrontal Cortex Schizophrenia Compared to Controls. Front Neurosci, 2022. 16: p. 858989.

65. Cai, H.Q., et al., Increased macrophages and changed brain endothelial cell gene expression in the frontal cortex of people with schizophrenia displaying inflammation. Mol Psychiatry, 2020. 25(4): p. 761–775.

66. Fernandes, B.S., et al., C-reactive protein is increased in schizophrenia but is not altered by antipsychotics: meta-analysis and implications. Mol Psychiatry, 2016. 21(4): p. 554–64.

67. Li, N., et al., Immunosenescence-related T cell phenotypes, structural brain imaging, and cognitive impairment in patients with schizophrenia: a moderated mediation analysis. Schizophrenia (Heidelb), 2025. 11(1): p. 101.

68. Chen, S., Y. Tan, and L. Tian, Immunophenotypes in psychosis: is it a premature inflamm-aging disorder? Mol Psychiatry, 2024. 29(9): p. 2834–2848.

69. Franceschi, C., et al., Inflammaging: a new immune-metabolic viewpoint for age-related diseases. Nat Rev Endocrinol, 2018. 14(10): p. 576–590.

70. Basurco, L., et al., Chronological versus immunological aging: Immune rejuvenation to arrest cognitive decline. Neuron, 2025. 113(1): p. 140–153.

71. Davila, C., et al., Perceptions of Patient-Clinician Communication Among Adults With and Without Serious Illness. JAMA Netw Open, 2025. 8(3): p. e250365.

72. Mazza, M.G., et al., Neutrophil-lymphocyte ratio, monocyte-lymphocyte ratio and platelet-lymphocyte ratio in non-affective psychosis: A meta-analysis and systematic review. World J Biol Psychiatry, 2020. 21(5): p. 326–338.

73. Maier, L.K., A. Marchfelder, and L. Randau, Meeting Report: German Genetics Society-Genome Editing with CRISPR. Bioessays, 2020. 42(2): p. e1900223.

74. Karageorgiou, V., G.P. Milas, and I. Michopoulos, Neutrophil-to-lymphocyte ratio in schizophrenia: A systematic review and meta-analysis. Schizophr Res, 2019. 206: p. 4–12.

75. Smetanova, J., et al., Accelerated Maturation, Exhaustion, and Senescence of T cells in 22q11.2 Deletion Syndrome. J Clin Immunol, 2022. 42(2): p. 274–285.

76. Mekori-Domachevsky, E., et al., Elevated Proinflammatory Markers in 22q11.2 Deletion Syndrome Are Associated With Psychosis and Cognitive Deficits. J Clin Psychiatry, 2017. 78(9): p. e1219–e1225.

77. Mekori-Domachevsky, E., et al., Neutrophils to lymphocytes ratio and psychosis in 22q11.2 deletion syndrome - Clinical and scientific implications. Schizophr Res, 2021. 231: p. 164–169.

78. Pascucci, A., et al., Developmental Alterations in the Diffusion Tensor Imaging Analysis Along the Perivascular Space Index Suggest Possible Glymphatic-Related Mechanisms Underlying Excitation/Inhibition Imbalance and Psychosis Vulnerability in 22q11.2 Deletion Syndrome. Biol Psychiatry Glob Open Sci, 2026. 6(3): p. 100713.

79. Hoang, D., et al., Inflammatory Subtypes in Antipsychotic-Najve First-Episode Schizophrenia are Associated with Altered Brain Morphology and Topological Organization. Brain Behav Immun, 2022. 100: p. 297–308.

80. Boerrigter, D., et al., Using blood cytokine measures to define high inflammatory biotype of schizophrenia and schizoaffective disorder. J Neuroinflammation, 2017. 14(1): p. 188.

81. Lu, H., L. Ai, and B. Zhang, TNF-α induces AQP4 overexpression in astrocytes through the NF-κB pathway causing cellular edema and apoptosis. Biosci Rep, 2022. 42(3).

82. Ito, H., et al., Interleukin-1beta induces the expression of aquaporin-4 through a nuclear factor-kappaB pathway in rat astrocytes. J Neurochem, 2006. 99(1): p. 107–18.

83. Li, Y., et al., Inhibition of astrocyte BMP signaling alleviates neuroinflammation in experimental models of Parkinson’s disease. Cell Death Discov, 2025. 11(1): p. 528.

84. Han, T., et al., Inflammatory stimulation of astrocytes affects the expression of miRNA-22-3p within NSCs-EVs regulating remyelination by targeting KDM3A. Stem Cell Res Ther, 2023. 14(1): p. 52.

85. Skauli, N., et al., Canonical Bone Morphogenetic Protein Signaling Regulates Expression of Aquaporin-4 and Its Anchoring Complex in Mouse Astrocytes. Front Cell Neurosci, 2022. 16: p. 878154.

86. Eixarch, H., et al., Inhibition of the BMP Signaling Pathway Ameliorated Established Clinical Symptoms of Experimental Autoimmune Encephalomyelitis. Neurotherapeutics, 2020. 17(4): p. 1988–2003.

87. Liu, Z., et al., Dysregulated Glial Differentiation in Schizophrenia May Be Relieved by Suppression of SMAD4- and REST-Dependent Signaling. Cell Rep, 2019. 27(13): p. 3832–3843.e6.

88. Xingi, E., et al., LPS-Induced Systemic Inflammation Affects the Dynamic Interactions of Astrocytes and Microglia with the Vasculature of the Mouse Brain Cortex. Cells, 2023. 12(10).

89. Yan, W., et al., β-Dystroglycan cleavage by matrix metalloproteinase-2/-9 disturbs aquaporin-4 polarization and influences brain edema in acute cerebral ischemia. Neuroscience, 2016. 326: p. 141–157.

90. Zhang, X., J. Zhang, and M. Shen, MitoQ attenuates maternal immune activation-induced mitochondrial and ER stress programs and ameliorates psychiatric-like behavioral deficits. Free Radic Biol Med, 2026. 253: p. 351–363.

91. Khandakar, R., et al., Prenatal and childhood infections as risk factors for psychiatric disorders: A systematic review and meta-analysis. Neurosci Biobehav Rev, 2026. 189: p. 106861.

92. Purcell, R.H., et al., Cross-species analysis identifies mitochondrial dysregulation as a functional consequence of the schizophrenia-associated 3q29 deletion. Sci Adv, 2023. 9(33): p. eadh0558.

93. Pihala, T., V. Kiviniemi, and M.H. Martikainen, Associations between the brain glymphatic system and mitochondrial (dys)function: a systematic review. Front Neurosci, 2025. 19: p. 1726054.

94. Eide, P.K., et al., Increased occurrence of pathological mitochondria in astrocytic perivascular endfoot processes and neurons of idiopathic intracranial hypertension. J Neurosci Res, 2021. 99(2): p. 467–480.

95. Hasan-Olive, M.M., et al., Pathological mitochondria in neurons and perivascular astrocytic endfeet of idiopathic normal pressure hydrocephalus patients. Fluids Barriers CNS, 2019. 16(1): p. 39.

96. Kopeć, K., et al., Glymphatic System and Mitochondrial Dysfunction as Two Crucial Players in Pathophysiology of Neurodegenerative Disorders. Int J Mol Sci, 2023. 24(12).

97. Kant, S., et al., Choroid plexus genes for CSF production and brain homeostasis are altered in Alzheimer’s disease. Fluids Barriers CNS, 2018. 15(1): p. 34.

98. Tsitsou-Kampeli, A., S. Suzzi, and M. Schwartz, The immune and metabolic milieu of the choroid plexus as a potential target in brain protection. Trends Neurosci, 2024. 47(8): p. 573–582.

99. Martini, H., et al., Mitochondrial metabolism and epigenetic crosstalk drive SASP. Nature, 2026. 656(8129): p. 980–992.

100. Grimm, A., U. Lang, and A. Eckert, Mitochondria and brain aging: From cell-specific dysfunction to intercellular cooperation. Neuron, 2026. 114(15): p. 2674–2690.

101. Varghese, N., et al., Tracing mitochondrial marks of neuronal aging in iPSCs-derived neurons and directly converted neurons. Commun Biol, 2025. 8(1): p. 723.

102. Hong, Y., et al., Reactive Oxygen Species Signaling and Oxidative Stress: Transcriptional Regulation and Evolution. Antioxidants (Basel), 2024. 13(3).

103. Pacca-Corrêa, J.B.L., et al., Astrocyte States in Brain Aging and Neurodegeneration: At the Crossroads of Senescence and Reactivity. Neurochem Res, 2026. 51(2).

104. Kim, D., et al., Senescent Astrocytes Derived from Human Pluripotent Stem Cells Reveal Age-Related Changes and Implications for Neurodegeneration. Aging Dis, 2024. 16(3): p. 1709–1731.

105. Diniz, L.P., et al., Accumulation of damaged mitochondria in aging astrocytes due to mitophagy dysfunction: Implications for susceptibility to mitochondrial stress. Biochim Biophys Acta Mol Basis Dis, 2024. 1870(8): p. 167470.

106. Wang, Y., et al., Mitochondrial-inflammation crosstalk in major depressive disorder: molecular mechanisms and therapeutic implications. Mol Psychiatry, 2026.

107. Santos-Silva, T., et al., Adolescent Stress-Induced Ventral Hippocampus Redox Dysregulation Underlies Behavioral Deficits and Excitatory/Inhibitory Imbalance Related to Schizophrenia. Schizophr Bull, 2025. 51(2): p. 501–512.

108. Gӧbel, J., et al., Mitochondria-Endoplasmic Reticulum Contacts in Reactive Astrocytes Promote Vascular Remodeling. Cell Metab, 2020. 31(4): p. 791–808.e8.

109. Thrane, A.S., et al., Critical role of aquaporin-4 (AQP4) in astrocytic Ca2+ signaling events elicited by cerebral edema. Proc Natl Acad Sci U S A, 2011. 108(2): p. 846–51.

110. Mlynek, E., et al., Disturbed Functional Connectivity Between Anterior Default Mode and Sensory Processing Regions Is Linked to Peripheral Inflammatory Markers and Psychopathology in Schizophrenia. Schizophr Bull, 2026. 52(1).

111. Leucht, S., et al., Dose Equivalents for Antipsychotic Drugs: The DDD Method. Schizophr Bull, 2016. 42 Suppl 1(Suppl 1): p. S90–4.

112. Keefe, R.S., et al., The Brief Assessment of Cognition in Schizophrenia: reliability, sensitivity, and comparison with a standard neurocognitive battery. Schizophr Res, 2004. 68(2-3): p. 283–97.

113. Gerstner, N., et al., Contrasting genetic predisposition and diagnosis in psychiatric disorders: A multi-omic single-nucleus analysis of the human OFC. Sci Adv, 2025. 11(10): p. eadq2290.

114. Gaebler, A.J., et al., Functional connectivity signatures of NMDAR dysfunction in schizophrenia-integrating findings from imaging genetics and pharmaco-fMRI. Transl Psychiatry, 2023. 13(1): p. 59.

115. Viher, P.V., et al., White matter correlates of the disorganized speech dimension in schizophrenia. Eur Arch Psychiatry Clin Neurosci, 2018. 268(1): p. 99–104.

116. Strik, W., et al., The Bern psychopathology scale for the assessment of system-specific psychotic symptoms. Neuropsychobiology, 2010. 61(4): p. 197–209.

117. Eisma, J.J., et al., Deep learning segmentation of the choroid plexus from structural magnetic resonance imaging (MRI): validation and normative ranges across the adult lifespan. Fluids Barriers CNS, 2024. 21(1): p. 21.

118. Smith, S.M., et al., Accurate, robust, and automated longitudinal and cross-sectional brain change analysis. Neuroimage, 2002. 17(1): p. 479–89.

119. Xie, K., et al., Association of immune-inflammatory biomarkers during pregnancy and the postpartum period with postpartum depression symptoms: A cross-sectional and longitudinal retrospective analysis. Brain Behav Immun, 2025. 129: p. 42–51.

120. Gong, W., et al., Glymphatic function and choroid plexus volume is associated with systemic inflammation and oxidative stress in major depressive disorder. Brain Behav Immun, 2025. 128: p. 266–275.

121. Durinck, S., et al., Mapping identifiers for the integration of genomic datasets with the R/Bioconductor package biomaRt. Nat Protoc, 2009. 4(8): p. 1184–91.

122. Wolf, F.A., P. Angerer, and F.J. Theis, SCANPY: large-scale single-cell gene expression data analysis. Genome Biol, 2018. 19(1): p. 15.

123. Korsunsky, I., et al., Fast, sensitive and accurate integration of single-cell data with Harmony. Nat Methods, 2019. 16(12): p. 1289–1296.

124. Traag, V.A., L. Waltman, and N.J. van Eck, From Louvain to Leiden: guaranteeing well-connected communities. Sci Rep, 2019. 9(1): p. 5233.

125. Hao, Y., et al., Dictionary learning for integrative, multimodal and scalable single-cell analysis. Nat Biotechnol, 2024. 42(2): p. 293–304.

126. Blondel, V.D., et al., Fast unfolding of communities in large networks. Journal of Statistical Mechanics: Theory and Experiment, 2008. 2008(10): p. P10008.

127. Leek, J.T., et al., The sva package for removing batch effects and other unwanted variation in high-throughput experiments. Bioinformatics, 2012. 28(6): p. 882–3.

128. Foroutan, M., et al., Single sample scoring of molecular phenotypes. BMC Bioinformatics, 2018. 19(1): p. 404.

129. Fox, J., M. Friendly, and S. Weisberg, Hypothesis tests for multivariate linear models using the car package. 2013.

130. Johnson, W.E., C. Li, and A. Rabinovic, Adjusting batch effects in microarray expression data using empirical Bayes methods. Biostatistics, 2007. 8(1): p. 118–27.

131. Gaser, C., et al., CAT: a computational anatomy toolbox for the analysis of structural MRI data. Gigascience, 2024. 13.

132. Ashburner, J. and K.J. Friston, Diffeomorphic registration using geodesic shooting and Gauss-Newton optimisation. Neuroimage, 2011. 55(3): p. 954–67.

133. Sled, J.G., A.P. Zijdenbos, and A.C. Evans, A nonparametric method for automatic correction of intensity nonuniformity in MRI data. IEEE Trans Med Imaging, 1998. 17(1): p. 87–97.

134. Ashburner, J. and K.J. Friston, Unified segmentation. Neuroimage, 2005. 26(3): p. 839–51.

135. Wang, H. and P.A. Yushkevich, Multi-atlas segmentation with joint label fusion and corrective learning-an open source implementation. Front Neuroinform, 2013. 7: p. 27.

136. de Lange, A.G. and J.H. Cole, Commentary: Correction procedures in brain-age prediction. Neuroimage Clin, 2020. 26: p. 102229.

137. Xu, Z., et al., methscore: a comprehensive R function for DNA methylation-based health predictors. Bioinformatics, 2024. 40(5).

138. Xu, Z., L. Niu, and J.A. Taylor, The ENmix DNA methylation analysis pipeline for Illumina BeadChip and comparisons with seven other preprocessing pipelines. Clin Epigenetics, 2021. 13(1): p. 216.

139. Higgins-Chen, A.T., et al., A computational solution for bolstering reliability of epigenetic clocks: Implications for clinical trials and longitudinal tracking. Nat Aging, 2022. 2(7): p. 644–661.

