## Supplementary table 1 for "Multimodal evidence for impaired glymphatic brain clearance as a mediator of “inflammaging” in schizophrenia – integrating insights from in vivo neuroimaging, omics, and post-mortem single nucleus-RNA-Seq data"

**Supplementary Table 1.** **Demographic data of SCZ and CTR groups**

| **Demographic data** | **SCZ** | **CTR** | **Statistical Test** | |
| --- | --- | --- | --- | --- |
|  |  |  | t(186)/χ^2^(1) | p |
| *In vivo cohort* | | | | |
| Number of subjects | n = 115 | n = 73 |  |  |
| Age(y) | 31.887 ±11.005 | 33.055 ±13.187 | 0.656 | 0.513 |
| Gender[male, n(%)] | 85(73.913%) | 40(54.795%) | 6.493 | 0.011 |
| HED [mg/d] | 7.551 ± 7.419 | -- |  |  |
|  |  |  | t(185)/χ^2^(1) | p |
| *Post-mortem cohort* | | | | |
| Number of subjects | n=95 | n=92 |  |  |
| Age(y) | 55.695 ±13.779 | 56.500 ±13.577 | 0.403 | 0.688 |
| Gender[male, n(%)] | 61(64.211%) | 56(60.870%) | 0.103 | 0.748 |

*Note:* CTR, control; HED, haloperidol equivalent dose; SCZ, Schizophrenia;
